# Structural basis of Ty3 retrotransposon integration at RNA Polymerase III-transcribed genes

**DOI:** 10.1101/2021.07.27.453949

**Authors:** Guillermo Abascal-Palacios, Laura Jochem, Carlos Pla-Prats, Fabienne Beuron, Alessandro Vannini

**Affiliations:** Division of Structural Biology, The Institute of Cancer Research, London SW7 3RP, United Kingdom; Friedrich Miescher Institute for Biomedical Research, Basel, Switzerland; Fondazione Human Technopole, Structural Biology Research Centre, 20157 Milan, Italy

## Abstract

Retrotransposons are endogenous elements that have the ability to mobilise their DNA and integrate at different locations in the host genome. In budding yeast, the Ty3 retrotransposon integrates with an exquisite specificity in a narrow window upstream of RNA Polymerase III-transcribed genes, such as the genes of transfer RNAs, representing a paradigm for specific targeted integration.

Here we present the cryo-EM reconstruction at 4.0 Å-resolution of an active Ty3 strand-transfer complex (Ty3 intasome) caught in the act of integrating onto a specific tRNA gene bound to the RNA Polymerase III general transcription factor TFIIIB, which is required for Ty3 specific targeting.

The structure unravels the molecular mechanisms underlying Ty3 integration specificity at RNA Polymerase III-transcribed genes and sheds light into the architecture of a retrotransposon integration machinery during the process of strand transfer at a genomic locus. The Ty3 intasome establishes contacts with a region of the TATA-binding protein (TBP), a subunit of TFIIIB, which is blocked by the ubiquitous transcription regulator negative cofactor 2 (NC2) in RNA Pol II-transcribed genes.

A previously unrecognised chromodomain of the Ty3 integrase mediates non-canonical interactions with TFIIIB and the tRNA gene itself, defining with extreme precision the position of the integration site. Surprisingly, Ty3 retrotransposon tethering to TFIIIB topologically resembles LEDGF/p75 transcription factor targeting by HIV retrovirus, highlighting mechanisms of convergent evolution by unrelated mobile elements and host organisms.

The Ty3 intasome-TFIIIB-tRNA promoter complex presented here represents a detailed molecular snapshot of a general transcription factor’s co-option by a mobile element, resulting in harmless integration into the host genome.

## MAIN

Transposable elements (TEs) are mobile DNA sequences that can affect integrity and stability of host genomes, potentially disrupting genetic information and providing sites for homologous recombination. Thus, not surprisingly, TEs are implicated in several human diseases^1, 2^. Long terminal repeat (LTR) retrotransposons are a class of TEs that constitute a significant fraction of eukaryotic genomes and their “copy-and-paste” mobilisation into new loci occurs via an RNA intermediate, relying on the intrinsic reverse transcriptase (RT) and integrase (IN) activities^3–5^.

Ty3/Gypsy is a member of the *Metaviridae* family of LTR-retrotransposons characterised by a highly defined targeting pattern in *S. cerevisiae*, integrating in a narrow location 2-3 bp upstream of RNA Pol III-transcribed genes^6, 7^. Ty3/Gypsy life cycle is similar to retroviruses but it is confined within a cell. Tethering of transposable elements and retroviruses to host factors, in order to integrate at specific genomic location, is not confined only to Ty3/Gypsy TEs^8–11^. For example, Ty1/Copia retrotransposon interacts directly with subunits of RNA Pol III^12–15^, *S. pombe* Tf1 element interacts with the Atf1p protein^8^ and the human immunodeficiency virus (HIV) binds to the transcriptional co-activator lens epithelium-derived growth factor (LEDGF/p75)^16–18^.

RNA Polymerase (Pol) III is responsible for the synthesis of short untranslated RNAs such as the U6 spliceosomal RNA or the transfer RNAs (tRNAs)^19^. The recruitment to its target genes depends on a set of general transcription factors (GTFs) that mediates very stable binding to specific DNA promoter elements^20^. TFIIIB, which is minimally required for initiation of Pol III transcription^21–25^, binds upstream of the transcription start site (TSS)^26^ and consists of the TATA-box binding protein (TBP), the B-related factor 1 (Brf1)^27^ and the B double prime (Bdp1) subunit^28^. TFIIIC, a 6-subunit GTF, recognises specific Pol III intragenic promoter elements and is required for the recruitment of TFIIIB at TATA-less promoters^29^.

The minimal factors required for Ty3 targeting at Pol III genes are the TFIIIB subunits, Brf1 and TBP, whereas TFIIIC is essential for directionality of integration, which occurs in a narrow window of 2-3 base pairs (bp) upstream the transcriptional start site (TSS)^30–32^. Direct interactions between TFIIIB and the Ty3 IN have been reported in the past but the molecular mechanisms by which such an exquisite specificity is achieved remain elusive. Ty3 represents a paradigmatic model to study retrotransposon’s targeted integration in host genomes. Notably, recent studies suggest that reactivation of endogenous retroelements in *H. sapiens*, a vertebrate endogenous retrovirus closely related to Ty3, is linked to disease development^33, 34^, including neurological disorders^35, 36^ and cancer^37, 38^. Despite recent advances^39, 40^, the structural analysis of host-retroelement targeting has been restricted to cryo-electron microscopy (cryo-EM) structures of histone-integrase interactions in nucleosomal context^41, 42^ and to crystal structures of HIV integrase domains bound to LEDGF/p75^16^. Here, we report the reconstitution of the Ty3 integration machinery (hereinafter referred as intasome), the biochemical characterisation of the Ty3 intasome interaction with the host factors TFIIIB and TFIIIC and, finally, the 4.0 Å-resolution cryo-EM reconstruction of a Ty3 intasome targeting and integrating at a TFIIIB-bound tRNA gene promoter.

### *In-vitro* reconstitution of an active Ty3 retrotransposon machinery

To obtain insights into the molecular mechanisms of Ty3 retrotransposition, we reconstituted the Ty3 intasome *in-vitro* using recombinant Ty3 integrase and Cy5- labelled oligonucleotides corresponding to the Ty3 gene LTR-termini (Fig. 1a and Extended Data Fig. 1a, b). In order to probe target-specific integration of the reconstituted Ty3 intasome, Ty3 activity was assessed by monitoring the specific recruitment to RNA Pol III transcription factors bound to a 157 bp-long synthetic *tD(GUC)K* gene promoter followed by the identification of the DNA products resulting from integration events (Fig. 1).

**Figure 1.**
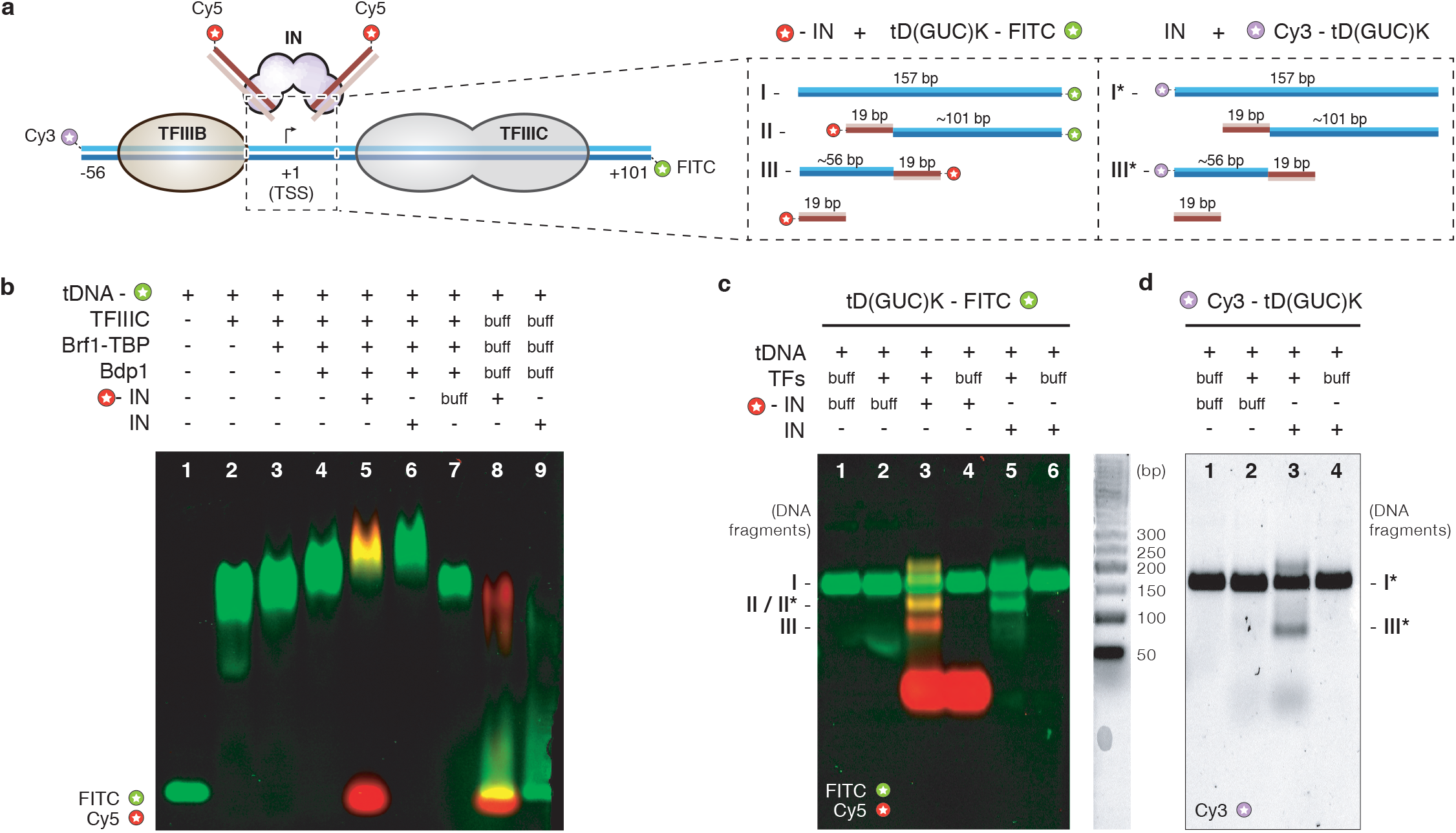
Assembly and activity of Ty3 retrotransposon machinery. **a,** Schematic representation of components used during binding and activity assays with reconstituted Ty3 retrotransposon machinery. *tD(GUC)K* template and non- template DNA strands are depicted as dark and light blue lines, respectively. DNA is numbered relative to the transcription start site (TSS). Reactive and non-reactive DNA strands of Ty3 LTRs are shown as dark and light red lines, respectively. TFIIIB and TFIIIC transcription factors, and intasome (IN) complex are depicted as globular shapes and coloured in wheat, grey and pink, respectively. Position and detail of the fluorophores used in the assays are shown as starred circles. The same colour scheme is used for fluorophores in **b-c**. *Inset,* summary and estimated length of the DNA products expected from integration events in the proximity of the TSS. **b,** Analysis of complex assembly. Electrophoretic mobility shift assay (EMSA) shows sequential binding of transcription factors to the *tD(GUC)K* DNA *(lanes 1-4)*, followed by binding of either Cy5-labelled Ty3 intasome (lane 5) or unlabelled intasome *(lane 6)* or intasome buffer *(lane 7)*. Incubation of Cy5-labelled Ty3 intasome *(lane 8)* or unlabelled intasome *(lane 9)* with *tD(GUC)K* template in the absence of transcription factors does not lead to a recruitment to the DNA. **c,** Integration activity assays of reconstituted Ty3 intasome (IN) into FITC-labelled *tD(GUC)K* tDNA or **d,** Cy3-labelled *tD(GUC)K* tDNA, in the presence or absence of pre-assembled TFIIIC and TFIIIB transcription factors (TFs). DNA products were analysed in 4.5% agarose gels after proteinase K treatment. Insets specify fluorophore detection (FITC, Cy3 or Cy5).

After step-wise binding of the TFIIIC transcription factor and TFIIIB to the *tD(GUC)K* gene promoter (Fig. 1b, lanes 2-4), a pre-assembled Ty3 intasome was added to the reaction. Electrophoretic mobility shift assays (EMSA) confirmed the ability of the reconstituted Ty3 intasome to specifically bind the *tD(GUC)K* gene promoter (Fig. 1b, lanes 5-6). The binding of the Ty3 intasome relies on the presence of the TFIIIB-TFIIIC general transcription factor complex as no binding was detected to the 157 bp FITC-labelled target DNA in their absence (Fig. 1b, lane 5 vs lane 8).

Evaluation of the integration activity was carried out following as a proxy the cleavage products of the *tD(GUC)K* nucleic acid scaffold, which would be expected from an integrative event by the *in vitro* Ty3 intasome, which is reconstituted with two linear short TR motifs as opposed to a single linear Ty3 gene with LTR termini (Fig. 1c, d). Taking into consideration that the TSS is asymmetrically located in the target DNA (Fig. 1a), predicted Ty3 integration 2-3 bp upstream the TSS would result in two DNA fragments (hereinafter, upstream and downstream fragments) of approximately 75 bp and 120 bp length, respectively (Fig. 1a, inset). To identify the DNA products, the integration reaction was performed on target DNAs labelled with FITC fluorophore at the downstream end and a Ty3 intasome assembled with Cy5-labelled LTRs (Fig. 1c). The presence of a *∼*120 bp band labelled with FITC and Cy5 confirmed the integration event and was only observed upon addition of Ty3 intasome (Fig. 1c, lane 2 vs lane 3, fragment II). To further verify the identity of this DNA fragment, the reaction was repeated using an unlabelled intasome, which provided a FITC-only band of a similar length (Fig. 1c, lane 5, fragment II*), confirming this as the downstream integration product. Similarly, a ∼75 bp Cy5-labelled band was also observed after Ty3 intasome addition, which was suggestive of an upstream integration product (Fig. 1c, lane 2 vs lane 3, fragment III). To confirm this hypothesis, a similar experiment was performed using a target DNA labelled only with a Cy3 fluorophore at the upstream end (Fig. 1d). This reaction produced a Cy3-labelled band of approximately 75 bp, which is only observed in the presence of Ty3 intasome (Fig. 1d, lane 2 vs lane 3, fragment III*), confirming this fragment as the upstream product of Ty3 integration. Reactions performed in the absence of TFIIIB and TFIIIC did not generate any integration products (Fig. 1c, lane 3 vs lane 4; and Fig. 1d, lane 3 vs lane 4), which confirmed the requirement of the transcription factors for Ty3 intasome recruitment to RNA Pol III-transcribed genes and the specificity of our *in vitro* assay. Lastly, analysis of the fluorescent products revealed also the presence of 157 bp non-integrated target DNA (Fig. 1c, fragment I; and Fig. 1d, fragment I*), which suggested that the *in vitro* reaction was not 100% efficient. Furthermore, in order to probe the role of TFIIIC in Ty3 integration, equivalent experiments were carried out in absence of TFIIIC (Extended Data Fig. 1d). In this context, TFIIIB alone was sufficient to efficiently recruit Ty3 intasome resulting in integration events qualitatively indistinguishable from the ones observed in presence of TFIIIC but with a reduced efficiency.

Overall, these results confirm that Ty3 *in vitro* reconstitution resulted in an active Ty3 retrotransposon machinery, displaying TFIIIB-dependent integration specificity that recapitulates Ty3 transposition at RNA Pol III-transcribed genes observed *in vivo*.

### Cryo-EM structure of Ty3 strand-transfer complex

To gain structural insights into the molecular mechanisms of Ty3 retrotransposon integration at RNA Pol III-transcribed genes, we purified the full and minimal integration machineries, assembled on TFIIIB-bound *tD(GUC)K* tRNA gene promoter in the presence or absence of TFIIIC transcription factor, respectively (Methods and Extended Data Fig. 1c, d).

Using the full integration machinery, we obtained a preliminary negative stain electron-microscopy (EM) reconstruction of the integration complex that showed the presence of an extra density not observed in TFIIIC-TFIIIB/DNA reconstructions, likely corresponding to the Ty3 intasome (Extended Data Fig. 1e). Further analysis by cryo-electron microscopy proved to be unsuccessful, likely due to conformational heterogeneity and low stability of the complex in the conditions used for sample freezing.

Analysis of the minimal integration complex by negative stain EM revealed highly detailed 2D class averages and provided a well resolved map where individual domains could be easily discerned and that was successfully used to fit TFIIIB and an *in-silico* model of Ty3 intasome (Extended Data Fig. 1f). We decided to further characterise this complex by cryo-EM. The un-crosslinked sample was applied to carbon-coated cryo-EM grids and data was collected at 0° and 30° tilting angles, to overcome issues with preferred particle orientation (Extended Data Fig. 2a, b). Following a process of hierarchical classification (Extended Data Fig. 2c, d), we obtained a cryo-EM reconstruction of the Ty3 intasome engaged with TFIIIB and the target DNA at an overall resolution of 4.0 Å (Extended Data Fig. 2e, f and Table 1).

**Table 1.** Cryo-EM data collection, refinement and validation statistics.

|  |  |
| --- | --- |
| **Data collection (0° Tilt)** |  |
| Voltage (kV) | 300 |
| Electron exposure (e<sup>-</sup>/Å<sup>2</sup>) | 52.1 |
| Defocus range (μm) | -1.7 to -3.2 |
| Pixel size (Å) | 1.047 |
| **Data collection (30° Tilt)** |  |
| Voltage (kV) | 300 |
| Electron exposure (e<sup>-</sup>/Å<sup>2</sup>) | 70.0 |
| Defocus range (μm) | -2.4 to -4.0 |
| Pixel size (Å) | 1.06 |
| **Reconstruction (RELION)** |  |
| Initial particle images (no.) | 628,998 |
| Final particle images (no.) | 101,469 |
| Map resolution (Å) | 3.98 |
| FSC threshold | (0.143-thr) |
| Map sharpening *B* factor (Å<sup>2</sup>) | -157.407 |
| **Model composition** |  |
| Non-hydrogen atoms | 21693 |
| Protein residues | 2250 |
| Nucleotide residues | 168 |
| **Refinement (PHENIX)** |  |
| Map CC (mask) | 0.59 |
| **R.m.s. deviations** |  |
| Bond lengths (Å) | 0.006 |
| Bond angles (°) | 1.113 |
| **Validation** |  |
| MolProbity score | 2.09 |
| Clashscore (all-atom) | 7.86 |
| Poor rotamers (%) | 0.00 |
| **Ramachandran plot** |  |
| Favored (%) | 84.77 |
| Allowed (%) | 15.10 |
| Disallowed (%) | 0.14 |

The cryo-EM map was used to build an atomic model of Ty3 retrotransposon minimal integration machinery (Extended Data Fig. 2g). The Ty3 intasome is a pseudo-symmetrical tetrameric complex, in which the four integrase subunits are engaged with two Ty3 long-terminal repeats (LTR). Two “core subunits” harbour the two active sites, placing in close juxtapositions the reactive LTR ends with the target DNA and are each associated with a peripheric subunit (Fig. 2). Sequence and structure comparisons show that the Ty3 retrotransposon is more closely related to the Prototype Foamy Virus (PFV)^43^ and *S. pombe* Tf1 retroelement than to other retroviruses such as the Rous sarcoma virus (RSV)^44^, the β-retrovirus mouse mammary tumour virus (MMTV)^45^ or the human immunodeficiency virus (HIV)^46^. Common to these mobile elements is a conserved core consisting of three domains: an HH-CC Zinc binding domain or N-terminal domain (NTD), a catalytic core domain (CCD), which adopts a RNAse H fold, and a SH3 domain or C-terminal domain (CTD) (Fig. 2b). Additionally, Ty3 encompasses a N-terminal extended domain (NED), which is involved in stabilising the interaction with the LTRs and has been reported to interact with TFIIIC^47^ (Fig. 2b and Fig. 3). Careful analysis of the structure reveals that a phosphodiester bond has been formed between the 3’-end of the LTR reactive strands and a cleaved 5’-end on the target DNA, confirming the activity of the assembled complex and suggesting that the majority of the particles imaged by cryo-EM correspond to a strand-transfer complex (Fig. 2b). The integration reaction takes place at position -6 (relative to the TSS) of the non-template strand and position -2 (relative to the TSS) of the template strand, resulting in a 5-bp target site duplication (TSD) (Fig. 3). The observed TSD and integration distance from the TSS recapitulate the previous finding observed *in vivo*^48^, supporting the existence of a preferred nucleotide sequence favouring Ty3 integration (Fig. 2b and Fig. 3).

**Figure 2.**
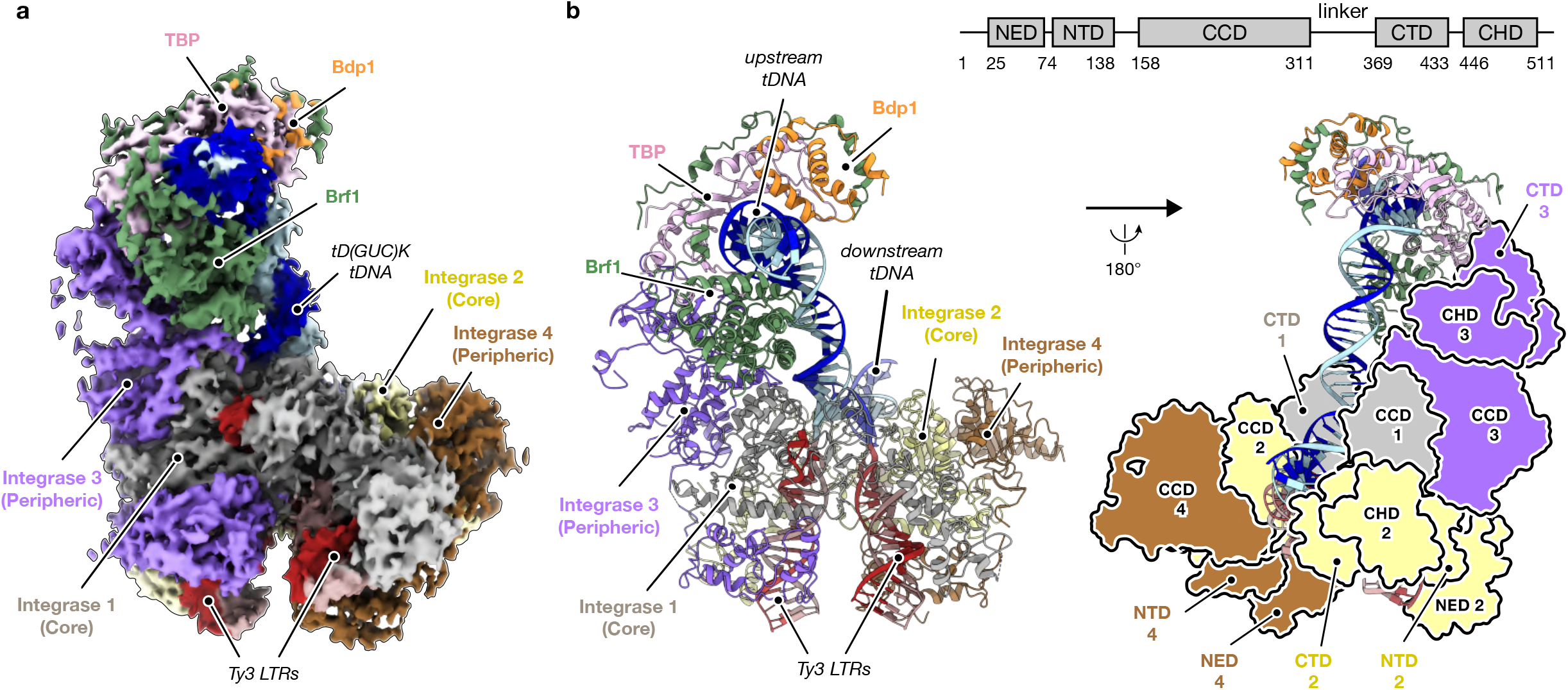
Cryo-EM structure of Ty3 strand-transfer complex bound to TFIIIB transcription factor and *tD(GUC)K* tRNA gene. **a,** Cryo-electron microscopy (cryo-EM) map of Ty3 retrotransposon strand-transfer complex (STC) bound to TFIIIB transcription factor and the t(GUC)K gene. Four integrase molecules are numbered and coloured in grey, yellow, purple and brown. Brf1, TBP and Bdp1 subunits of TFIIIB are depicted in green, pink and orange, respectively. Template and non-template strands of the tDNA are shown in dark and light blue, respectively. Ty3 long-terminal repeats (LTRs) are highlighted in red shades. **b,** Ribbon representation (front view, *left*) of the structure model coloured and numbered as described in **a** and back view (*right*) of Ty3 retrotransposon model representing as silhouettes the domain architecture of Ty3 integrase subunits (*inset*). NED, N-terminal extended domain; NTD, N-terminal domain; CCD, catalytic core domain; CTD, C-terminal domain; CHD, chromodomain.

**Figure 3.**
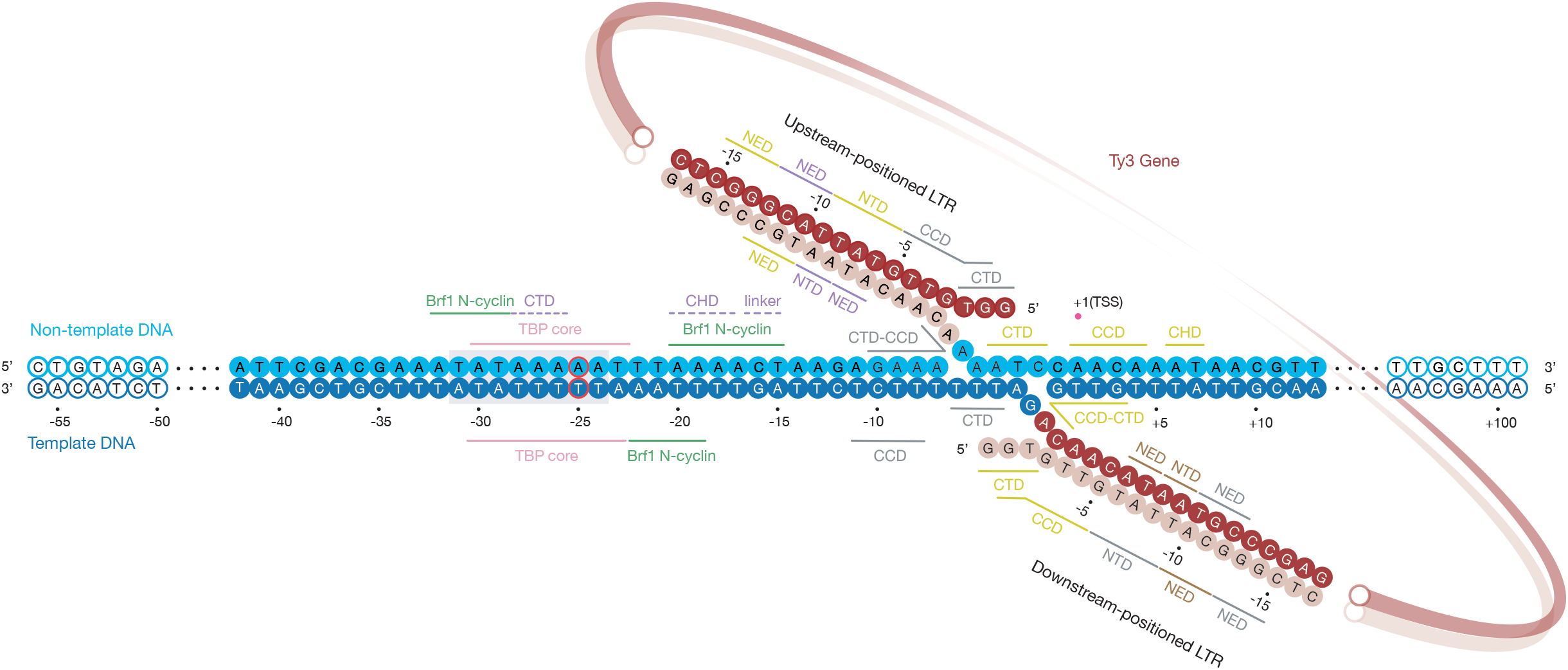
Details of protein-DNA interactions and integration site location on RNA Pol III-transcribed genes. DNA nucleotides of the *tD(GUC)K* gene modelled in the Ty3 strand-transfer complex are depicted as solid circles and numbered relative to the TSS. Template and non- template strands of the tDNA are shown in dark and light blue, respectively. Mutation of nucleotide -25 to break the pseudo-symmetry of the TATA box is outlined in red. The TATA box is highlighted with a grey box. Ty3 long-terminal repeats (LTRs) are highlighted in red shades. Protein-DNA and protein-protein interactions are indicated with solid and dashed lines, respectively.

Ty3 intasome engagement with the target DNA causes a ∼90° bent near the transcription start site, which is still compatible with binding to the TFIIIB transcription factor, in agreement with recent barcoding experiments highlighting the bendability of this region as an important determinant of frequency of integration^48^ (Fig. 2b). Model building of TFIIIB on the cryo-EM map led to the identification of the TBP core and Brf1 B-core cyclin-repeats. However, given the poor resolution around the Bdp1 transcription factor, only a rigid-body fit of the SANT domain could be carried out, (Fig. 2b and Extended Data Fig. 2g). Interaction of TFIIIB with the Ty3 intasome is mainly mediated by the C-terminal region of a peripheral Ty3 integrase subunit, which stablishes a novel set of contacts with the convex surface of TBP stirrups and Brf1 cyclin-repeats (Fig. 2b). The binding of TFIIIB far upstream (-15 to -32 bp) the TSS constrains the positioning of Ty3 intasome active site also upstream of the TSS, ensuring that RNA Pol III-transcribed genes are not disrupted by the integration event.

### Determinants of target specificity at RNA Pol III-transcribed genes

Initial model building of the intasome-TFIIIB interface revealed the involvement of the Ty3 integrase C-terminal domain (CTD) (Fig. 4a). The SH3 fold (residues 369-433) of the CTD of a peripheric Ty3 subunit establishes direct protein-protein interactions with the H2’ helix of TBP C-terminal lobe, positioning the Ty3 intasome upstream the TSS (Fig. 4a). Remarkably, the same region of Ty3 IN, characterised by a compact *β*-barrel architecture, also mediates tight interactions with the target DNA minor groove when in the context of the core Ty3 subunits, highlighting the plasticity of Ty3 domains in mediating different functions (Fig. 3). Topologically, this TBP-Ty3 interface overlaps with the binding interface of TBP and the RNA Pol II-specific transcription regulator negative cofactor 2 (NC2)^49^ (Fig. 4b). Thus, NC2*β* subunit binding to this region of TBP prevents recruitment of Ty3 intasome to RNA Pol II-transcribed genes before formation of a fully functional pre-initiation complex, rationalising the absence of integration at protein coding genes.

**Figure 4.**
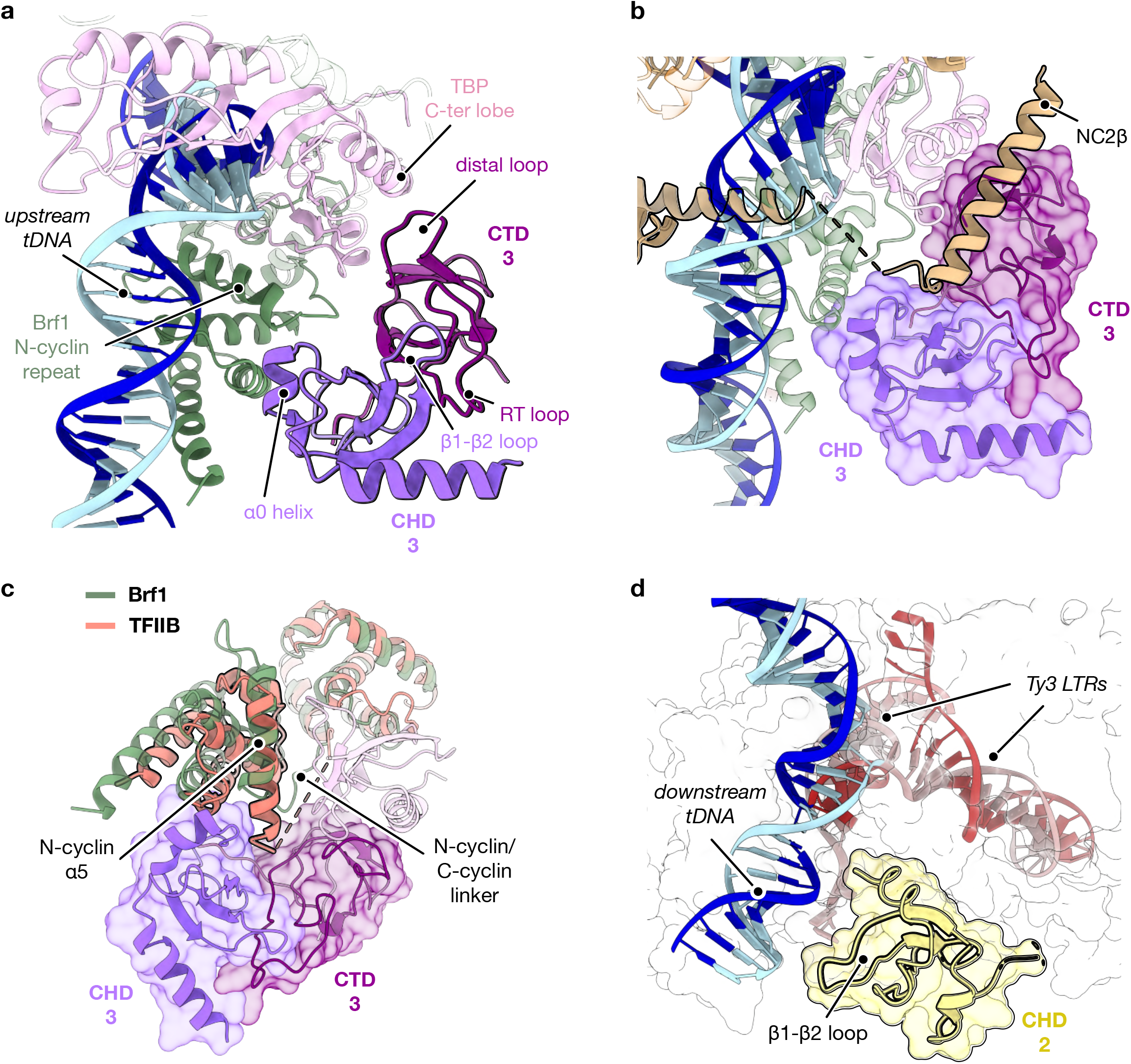
Architecture of TFIIIB – Ty3 interaction and molecular basis of integration specificity. **a,** Detail of TFIIIB transcription factor and Ty3 integrase binding domains, represented as ribbon models. The distal loop of Ty3 integrase C-terminal domain (CTD 3, purple) interacts with TBP C-terminal lobe (pink) whereas Ty3 integrase vestigial chromodomain (CHD 3, violet purple) is recruited to Brf1 N-terminal cyclin repeat (green) through its helix 0 (a0). CTD 3 and CHD 3 domains associate through the RT loop and the {31-{32 loop, respectively, adopting a novel tandem small {3-barrel conformation. **b,** Binding of the transcription regulator negative cofactor 2 (NC2{3, wheat ribbon, pdb code: 4WZS) and Ty3 intasome to TBP C-terminal lobe requires the same interface, which compromises the recruitment of Ty3 retrotransposon (purple, molecular surface) to RNA Pol II transcriptional units. **c,** The B-core cyclin repeat architecture of TFIIB (orange) and Brf1 (green) transcription factors differs in the length of the N-cyclin helix 5 (a5) and the N/C cyclin linker, which prevents binding of Ty3 intasome (purple, molecular surface) to TFIIB and integration at RNA Pol II- transcribed genes. **d,** The vestigial chromodomain of a symmetrically-related Ty3 integrase subunit (CHD 2, yellow) establishes contacts with the target DNA (tDNA) downstream of Ty3 LTRs integration site.

Further inspection of our cryo-EM map revealed the presence of an additional unidentified density close to the peripheral Ty3 CTD and TFIIIB. *De novo* model building allowed us to identify this region as a previously uncharacterised chromodomain (CHD), which was thought to be absent in members of the Ty3/Gypsy family in *S. cerevisiae*^50^ (Fig. 4a). This domain (residues 446-511) is located C-terminally to the CTD, forming together a novel compact tandem small *β*-barrel organisation^51^, which leads to a tighter binding to the TFIIIB transcription factor (Fig. 4a). Ty3 CHD shares the canonical motif organisation (*β*_1_-L_1_-*β*_2_-L_2_-*β*_3_-L_3_-*α*_1_-*α*_2_) but it lacks the conserved aromatic cage that is required for histone recognition in similar domains^52^ (Extended Data Fig. 3a), likely explaining why it remained unnoticed. Notably, the Ty3 CHD displays an additional N-terminal *α*-helix (*α*_0_; aa. 448-456), which folds back into the domain and partially mimics the binding of histone-tail peptides^53–55^ (Extended Data Fig. 3b-d). In the context of TFIIIB binding, the *α*_0_ helix acts as a “staple” that mediates interactions between the Ty3 integrase and the Brf1 N-terminal cyclin repeat (Fig. 4a). In particular, Ty3 CHD contacts the linker region between Brf1 cyclin repeats, which presents significant differences with TFIIB, the RNA Pol II orthologue of Brf1 (Fig. 4c and Extended Data Fig. 4a). In *S. cerevisiae,* this region of TFIIB includes an unstructured extension which is absent in Brf1 (Extended Data Fig. 4a) and that could represent a hindrance to Ty3 intasome recruitment, rationalising the absence of Ty3 integration at RNA Pol II-transcribed genes.

In our cryo-EM map, additional unattributed density was observed at the downstream end of the target DNA (Fig. 2a and Fig. 4d). Despite the poorer quality of the map in this region, the density could be unequivocally identified as the CHD of a core Ty3 integrase subunit. In this context, the Ty3 CHD mediates interactions with the *tD(GUC)K* gene downstream of the transcription start site (Fig. 3 and Fig. 4d). Thus, the positioning of CHDs stemming from two distinct Ty3 subunits at opposite ends of the integration site plays a major role in defining the docking of Ty3 intasome between the TATA-box and the TSS, ensuring that the integration event does not disrupt gene-body regions.

Highlighting the paramount role of CHDs for efficient Ty3 retrotransposition, *YILWTy3- 1*, one of two full-length Ty3 genes in *S. cerevisiae*, encompasses a frameshift mutation that disrupts the CHD, resulting in inactivation of this Ty3 retrotransposon that could be re-activated after restoration of the protein reading frame^56^.

Analogously to Ty3, CHDs also play a fundamental role in *S. pombe* Tf1 retrotransposon integration, where a non-canonical CHD (Extended Data Fig. 3a) promotes binding to the target DNA and it is required for integration into intergenic regions^8, 57, 58^, pointing towards a more general role of CHDs for efficient transposition of Ty3/Gypsy retroelements across different species.

### Ty3 intasome recruitment by TFIIIB requires an extended CCD-CTD linker

Recruitment of Ty3 intasome to RNA Pol III-transcribed genes depends on the interaction between TFIIIB and Ty3 integrase C-terminal region, which questions whether other CTD-containing retroelements could exploit similar mechanisms to target transcription factors and/or DNA bound complexes. Among the conserved integrase core domains, the main difference between Ty3 and other integrases such as RSV, MMTV or HIV, lays on the presence of an extended CCD-CTD linker, which in Ty3 folds into a long *α*-helix (*α*_10_) (Fig. 5a and Extended Data Fig. 4b). This feature is especially significant in the context of TFIIIB targeting because the *α*_10_ linker allows the positioning of Ty3 CTD and CHD at the outer boundaries of the intasome, acting as a platform for TFIIIB anchoring (Fig. 5a). Due to the reduced length of the CTD-CCD linker in RSV, MMTV and HIV retroviruses, their CTDs are positioned close to the intasome core, which prevents the interaction with factors positioned away from the body of the intasome (Fig. 5a). Conversely, the PFV retrovirus CTD-CCD linker displays a similar length and adopts a similar orientation compared to Ty3 integrase, suggesting targeting of more distant factors (Fig. 5a and Extended Data Fig. 4b).

**Figure 5.**
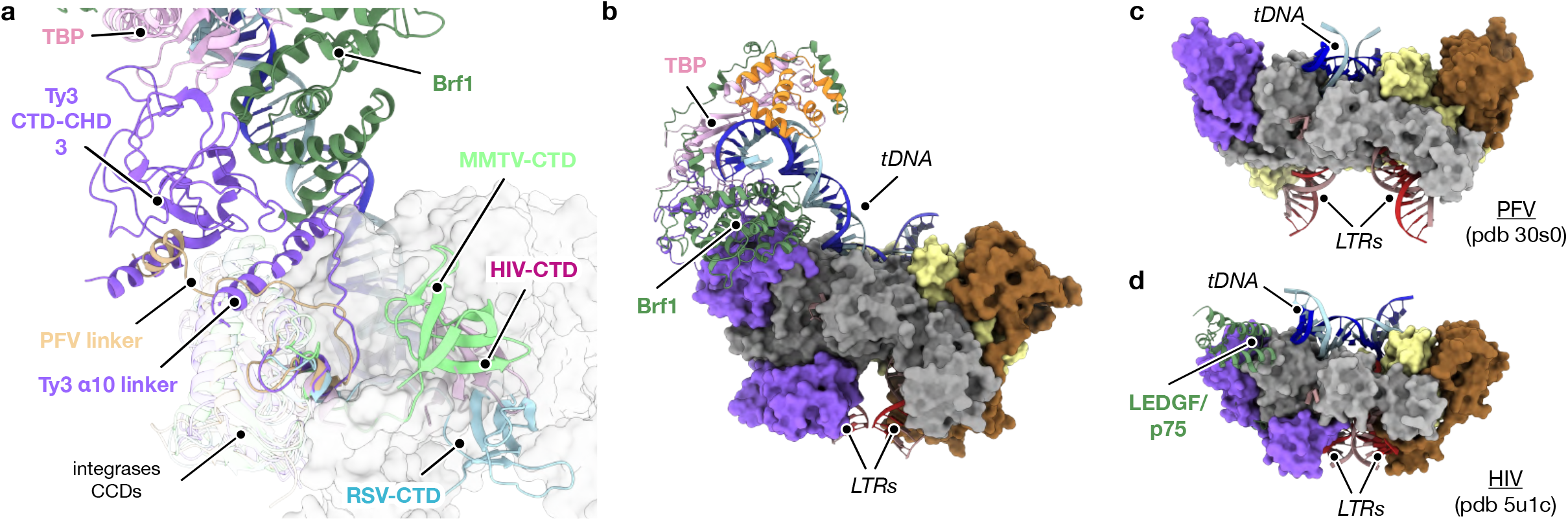
Ty3 CCD-CTD extended linker creates a platform for TFIIIB assembly. **a,** Detailed view of Ty3 integrase extended CCD-CTD linker conformation. Retroviral intasome models of PFV (pdb code: 3OS0, wheat), HIV (pdb code: 5U1C, pink), MMTV (pdb code: 3JCA, bright green) and RSV (pdb code: 5EJK, light blue) were superposed to the external catalytic core domain (CCD) of Ty3 integrase (purple). While retroviral CTDs are positioned close to the integrase core, Ty3 extended α10 linker adopts a peripheral orientation that creates a platform for the interaction between Ty3 CTD-CHD and TFIIIB transcription factor. The core of the intasome is shown as grey molecular surface. Structure comparison of **b,** Ty3 integrase with **c,** PFV and **d,** HIV integrases. Integrase core is shown as molecular surface and coloured as in Fig. 2. The general intasome architecture is conserved between the retrotransposon and retroviral integrases. Binding of LEDGF/p75 transcription factor to HIV is mediated by a similar region than in the Brf1-IN interaction, which highlights a potential hot-spot for host-integrase recognition events.

The overall architecture of the Ty3 intasome (Fig. 5b) resembles the organisation of other retroelements such as PFV or HIV (Fig. 5c, d). Intriguingly, in spite of the differences in the peripheral integrase subunits, the binding of the lens epithelium-derived growth factor (LEDGF/p75) to the HIV intasome is topologically related to the region that mediates the TFIIIB-Ty3 interaction (Fig. 5d). Similar to Brf1 B-core cyclin- repeats, the integrase binding domain of LEDGF/p75 factor also adopts a helical bundle organisation (PDB codes: 5U1C and <u>2B4J</u>)^16, 46^. This factor is required for HIV targeting host genomes and its depletion prevents retroviral integration and replication^17, 59–61^. Despite the distant link between both retroelements, the use of similar interfaces hints at the existence of “hot-spot” regions in the intasome periphery which are exploited to targeted DNA-binding factors in the host organism.

### Ty3 integration and RNA Pol III transcription are mutually exclusive

The central role of TFIIIB factor in Pol III recruitment and transcription initiation has been extensively characterised^21–25, 62^ but it remains to be understood how this is affected by other TFIIIB-mediated processes occurring at the same loci, such as Ty3 retrotransposon integration. The cryo-EM structure of TFIIIB targeted by a Ty3 intasome allowed us to compare these two unrelated processes. Superimposition of Ty3-TFIIIB complex with that of the Pol III pre-initiation complex (PIC)^21^ (Fig. 6) reveals that the architecture of TFIIIB is unaltered, except for the destabilisation of Bdp1 binding in the retrotransposon targeting complex reconstituted *in vitro*, which is essential for DNA opening during transcription initiation. Furthermore, the comparison of the two structures reveals that the Ty3 intasome and Pol III would occupy nearly the exact same position, thus providing mechanistic evidence that Pol III transcription and Ty3 integration are mutually exclusive, as previously postulated by studies suggesting competition between the two processes^48, 63^. Unlike Ty3, Ty1 retrotransposon integrates at nucleosomal DNA and requires a physical association with the Pol III enzyme^13, 14, 64^ rationalising why Ty3 and Ty1 do not compete for the same integration sites.

**Figure 6.**
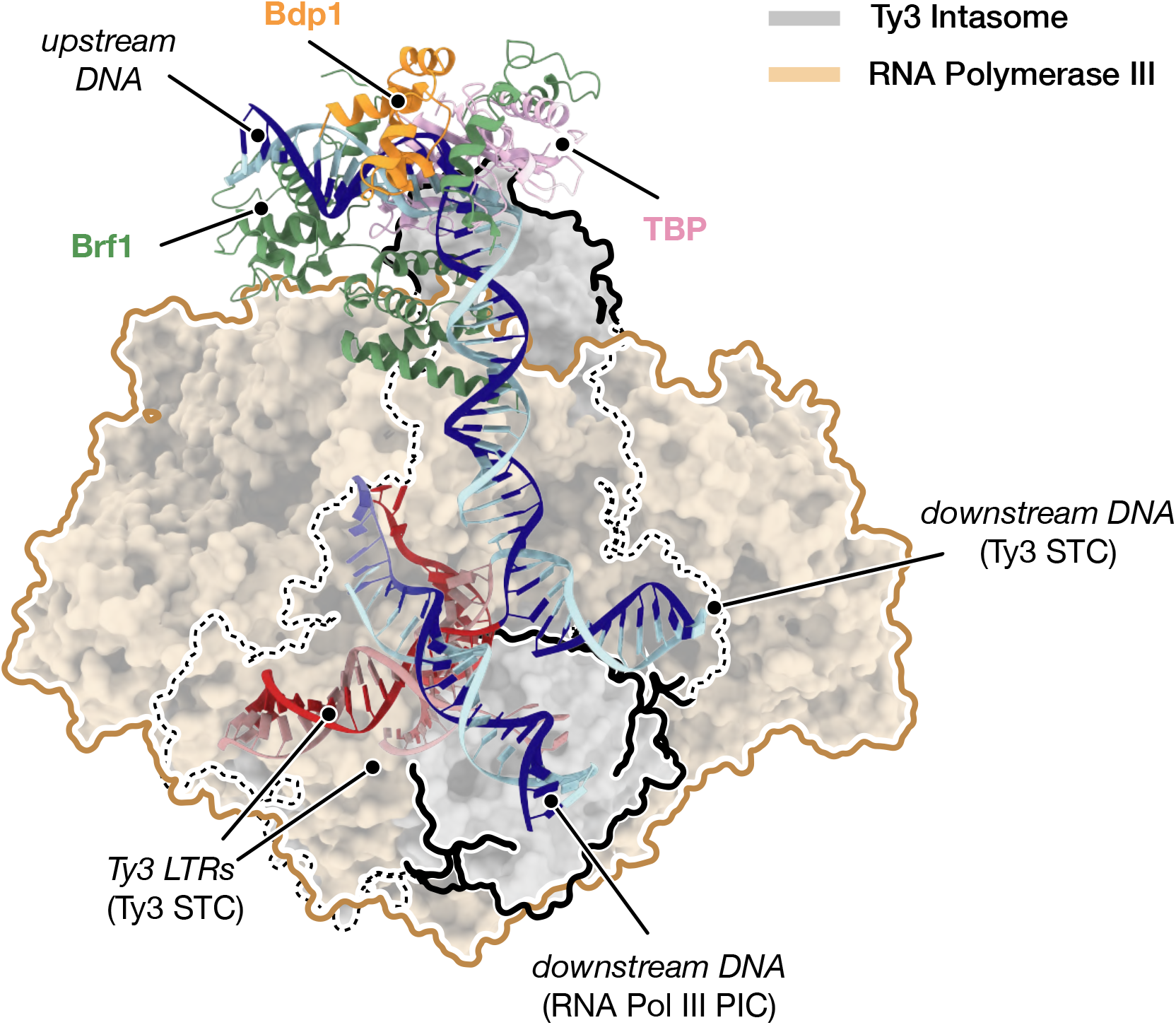
General comparison of TFIIIB targeting by RNA Polymerase III and Ty3 intasome. Comparison of RNA Polymerase III and Ty3 intasome recruitment by TFIIIB transcription factor. TFIIIB subunits are depicted as ribbons and coloured as described in Fig. 2. *S. cerevisiae* models of RNA Pol III pre-initiation complex (RNA Pol III PIC, pdb code: 6EU0) and Ty3 strand-transfer complex bound to TFIIIB (Ty3 STC, this work) were superimposed onto TBP subunit of TFIIIB. RNA Pol III and Ty3 intasome are depicted as wheat and grey molecular surfaces, and their boundaries are highlighted with brown and black lines, respectively. Binding of both enzymatic machineries to RNA Pol III-transcribed genes requires equivalent regions, which prevents simultaneous RNA transcription and integration events. Models of the target DNA are shown as ribbon and coloured as described in Fig. 2. Whereas the orientation of the upstream DNA is conserved, the direction of the tDNA in the proximity of the TSS differs upon binding of the two enzymes.

## DISCUSSION

Retrotransposons are the main active TEs in eukaryotes and are structurally and functionally related to retroviruses. The selection of integration sites clearly impacts on the consequences of mobilisation of TE elements on the host biology. Thus, insertions in deleterious locations are negatively selected. In this context, the Ty3 family of TE has evolved exploiting the peculiar architecture of Pol III-transcribed genes, which are defined by ubiquitous internal promoter elements such as the A and B boxes, which are binding sites of transcription factor TFIIIC^65–67^. By tightly binding to upstream transcription factor TFIIIB, which is ultimately recruited by TFIIIC, the Ty3 intasome drives the insertion of new copies of the Ty3 gene upstream of the transcribed region, while concomitantly leaving the promoter elements untouched. This strategy confers Ty3 retrotransposon the ability to integrate into safe-haven locations of the host genome while minimising deleterious effects.

Our cryo-EM reconstruction of Ty3 intasome bound to a tRNA gene promoter recapitulates previous biochemical data and integrates them into a model that involves the formation of an extensive network of interactions between TFIIIB transcription factor and a peripheral Ty3 integrase subunit. The structural data unveils how a newly-discovered chromodomain in Ty3 integrase mediates histone-independent interactions with Brf1 and the downstream part of the target DNA, acting as a “ruler” that safeguards the integration upstream of the transcription start site and prevents gene disruption (Fig. 3 and Fig. 4). Our atomic model deciphers the molecular determinants of target specificity at RNA Pol III-transcribed genes, highlighting how Ty3 intasome recruitment requires a region in the Brf1-TBP interface that is blocked by the NC2*β* factor in the TFIIB homologous factor at RNA Pol II-transcribed genes (Fig. 4). The incompatibility between Pol III machinery activity and Ty3 integration additionally allows for these two processes to be timely separated avoiding a direct overlap that could result in dangerous loss of genetic information.

Notably, our results found that Brf1 targeting by Ty3 element topologically resembles the recruitment of HIV to LEDGF/p75 factor, suggesting that host recognition occurs through similar mechanisms and interfaces across distant retroelements.

## METHODS

### Protein expression and purification

A pOPINF plasmid containing *S. cerevisiae* full-length Ty3 integrase was transformed into *E. coli* BL21-RIL-CodonPlus competent cells and grown at 37 °C to OD_600_ of ∼1 in LB medium supplemented with 50 μM ZnCl_2_. The cell culture was stored at 4 °C for 1h and an overnight induction was performed with 1 mM IPTG at 20 °C. Following cell harvest, the collected pellet was suspended in lysis buffer containing 20 mM HEPES pH 8, 750 mM NaCl, 10 mM MgCl_2_, 10 μM ZnCl_2_, 10 mM β-mercaptoethanol, 10 mM imidazole, 50 mM L-arginine and 50 mM L-glutamic acid, supplemented with two protease inhibitor tablets (Roche) and a scoop of DNAse, and adjusted to pH 7.4. After incubation for 30 min at 4 °C, cells were subjected to sonication (12 cycles, 15 s ON, 59 s OFF, 60% amplitude) and the lysate was fractionated by centrifugation at 48,000g for 40 min at 4°C. Then, the soluble fraction was filtered and incubated for 3 h at 4 °C with ∼6 ml of HisPur^TM^ Ni-NTA resin (Thermo Fisher Scientific). The beads were subsequently washed with ∼400 ml lysis buffer (without protease inhibitors or Arginine/Glutamic acid) and the protein eluted in 20 ml lysis buffer supplemented with 500 mM imidazole. N-terminal His-tag was removed by overnight incubation with ∼1 mg of 3C protease at 4 °C. The cleaved sample was then diluted to 125 mM NaCl with buffer HepA (20 mM HEPES pH 8, 10 mM MgCl_2_, 10 μM ZnCl_2_ and 1 mM DTT) and loaded into a HiTrap Heparin HP 5 ml column (GE Healthcare) equilibrated with 6.25% of buffer HepB (buffer HepA supplemented with 2 M NaCl). After washing, sample elution was performed through a linear gradient from 6.25% to 100% of buffer HepB in 30 CV. Fractions corresponding to the elution peak were analysed by SDS-PAGE and those containing Ty3 integrase were pooled, concentrated to ∼4 ml and loaded into a HiLoad 16/600 Superdex 200 pg column (GE Heathcare) equilibrated with 20 mM HEPES pH 8, 730 mM NaCl, 10 mM MgCl_2_, 10 μM ZnCl_2_ and 1 mM DTT. The fractions corresponding to pure Ty3 integrase were pooled, concentrated to ∼3.25 mg/ml, flash-frozen in liquid nitrogen and stored at -80 °C. The final yield was ∼2 mg of protein per litre of culture.

The six-subunit TFIIIC transcription factor was cloned into pBIG2ab vector using the biGBac system^68^. After plasmid transformation into the DH10EmBacY strain using the heat-shock technique, the cells were plated on an agar plate containing X-Gal, gentamycin, kanamycin and tetracycline and incubated overnight at 37 °C. White colonies corresponding to successful plasmid transposition were selected, expanded in 5 ml of LB media and the bacmid containing the TFIIIIC genes was purified. The generated plasmid was then mixed with Cellfectin^TM^ II reagent (Thermo Fisher Scientific) in a 1:1 molar ratio and transfected into adherent Sf9 insect cells at a density of 5 × 10^5^ cells per millilitre. Following incubation at 27 °C for 72 h, the cell culture supernatant (∼2 ml), which contained the P1 viral fraction, was collected and mixed with 25 ml of suspended Sf9 cells at a density of 5 × 10^5^ cells per millilitre. Cell viability and visualisation of the yellow fluorescent protein (YFP) in the cells for 3-5 days were used as an indicator of viral infection. The supernatant corresponding to the P2 viral fraction was collected once fluorescence was observed in 80-90% of the cells. Finally, protein expression was achieved by the addition of 2 ml of P2 fraction to 500 ml of suspended High5 insect cells at a density of 5 × 10^5^ cells/ml and growth for 3-4 days at 27 °C in Lonza Insect Xpress media. Cells were harvested and the pellet stored at -80 °C.

The insect cells pellet was suspended in ∼125 ml of lysis buffer containing 20 mM HEPES pH 8, 500 mM NaCl, 1 mM MgCl_2_, 10% glycerol and 10 mM β-mercaptoethanol, supplemented with a scoop of DNase I, two protease inhibitor tablets (Thermo Fisher Scientific) and 4 μl of benzonase. After incubation, the sample was subjected to sonication (9 cycles, 5 s ON, 10 s OFF, 20% amplitude) and fractionated by centrifugation at 37,565g for 30 min at 4°C. The soluble fraction was filtered and loaded into a StrepTrap HP 5ml column (GE Healthcare) equilibrated with lysis buffer. Following a wash step, the bound sample was eluted in 15 ml of lysis buffer supplemented with 0.05% (w/v) D-desthiobiotin. The sample was diluted to ∼150 mM NaCl with buffer HepA (20 mM HEPES pH 8, 10% glycerol and 10 mM β-mercaptoethanol) and loaded into a HiTrap Heparin HP 5ml column equilibrated with 7.5% of buffer HepB (buffer HepA supplemented with 2 M NaCl). After washing, elution from the column was performed through a linear gradient from 7.5% to 100% of buffer HepB in 10 CV. The fractions corresponding to the elution peak were analysed by SDS-PAGE and those containing the TFIIIC complex were collected and loaded into a XK 16/70 Superose 6 pg column (GE Healthcare) equilibrated with 20 mM HEPES pH 8, 200 mM NaCl, 2.5% glycerol and 1 mM DTT. Pure TFIIIC fractions were pooled, concentrated to ∼ 12.5 mg/ml, flash-frozen and stored at -80 °C.

Purifications of *S. cerevisiae* Brf1-TBP fusion protein and Bdp1 transcription factor were performed as described elsewhere^21^.

### *tD(GUC)K* target DNA

Fluorescently-labelled DNA fragments used for *in vitro* binding and integration assays were generated by large-scale PCR amplification of the *tD(GUC)K* promoter between -56 and +101 (according to TSS position) using *Pyrococcus abysii* DNA polymerase (PabPolB) purified in house. The generated DNA was precipitated with ice cold 100% ethanol (v/v), suspended in buffer QA (10 mM Tris pH 8 and 1 mM EDTA) and loaded into a MonoQ 5/50 GL column (GE Healthcare). After a step-wise wash with 10% and 20% of buffer QB (Buffer QA supplemented with 2M NaCl), elution was performed through a linear gradient from 20% to 50% of buffer QB in 40 CV. After analysis in 2% agarose gels, fractions containing pure DNA were pooled, precipitated with ice cold 100% ethanol and suspended in TE buffer pH 8 to a final concentration of ∼70 μM. The resulting 157-bp DNA fragment included the TATA-box, and the A- and B-box elements required for TFIIIC binding. Labelling of the upstream or downstream termini with Cy3 or FITC, respectively, was achieved through incorporation of the fluorophores in the amplification oligonucleotides.

Target DNA oligonucleotides used for cryo-EM analysis of Ty3 intasome binding to TFIIIB transcription factor included the TATA box and encompassed nucleotides -42 to +13 (according to TSS position) of the yeast *tD(GUC)K* gene promoter (template strand: 5’- AACGTTATTTGTTGGATTTTTTCTCTTAGTTTTAAATTTTTATATTTCGT CGAAT -3’ and non-template strand: 5’- ATTCGACGAAATATAAAAATTTAAAACTAAGAGAAAAAATCCAACAAATAACGTT -3’; Integrated DNA Technologies). As described previously^69^, a single nucleotide mutation in the TATA-box element was included in order to favour unidirectional positioning of TFIIIB.

### Ty3 intasome complex formation

The Ty3 intasome assembly was carried out using a 5’-overhang fragment of the retrotransposon LTR terminus, which consisted of a 17-bp reactive strand (5’-GAGC CCGTAATACAACA-3’; IDT) and a 19-bp non-reactive strand (5’-GGTGTTGTATT ACGGGCTC-3’; IDT). For *in vitro* binding and activity assays, a Cy5 fluorophore at the 5’ end of the reactive strand was included. The oligonucleotides were suspended and annealed as previously described^21^. The intasome assembly was carried out using 1.25 nmol of annealed LTRs and a 2.4-fold excess of pure Ty3 integrase, in 425 μl of buffer containing 20 mM HEPES pH 8, 730 mM NaCl, 10 mM MgCl_2_, 10 μM ZnCl_2_ and 1 mM DTT. The mixture was subjected to a progressive reduction of the salt concentration through overnight dialysis in 20 mM HEPES pH 8, 200 mM NaCl, 25 μM ZnCl_2_ and 2 mM DTT, using a 1 kDa MWCO Mini Dialysis Kit (GE Healthcare). Finally, the concentration of NaCl in the sample was increased to 320 mM by addition of NaCl 5M. Intasome formation was assessed by gel-filtration chromatography (Extended Data Fig. 1a) followed by SDS-PAGE analysis and detection of the Cy5 fluorophore in a native agarose gel (Extended Data Fig. 1b).

### Assembly of full and minimal integration machinery

*In vitro* targeting of the Ty3 intasome at the full RNA Pol III transcriptional machinery (i.e. containing TFIIIC and TFIIIB) was achieved through a process of sequential binding of transcription factors and integration machinery to the target DNA. First, 200 μg of TFIIIC transcription factor and the annealed 157-bp *tD(GUC)K* DNA were mixed in a 2.2:1 molar ratio and incubated at RT for 30 min (Fig. 1b, lane 1). Next, a 2.5-fold excess (relative to the DNA) of Brf1-TBP fusion protein and Bdp1 were added to the mixture in two consecutive steps and subjected to a similar incubation protocol (Fig. 1b, lanes 2-3). Finally, the reaction was supplemented with 400 μl of Cy5-labelled Ty3 intasome (assembled as described before) and the targeting allowed for 30 min at RT (Fig. 1b, lane 4). Controls were performed with un-labelled Ty3 intasome (Fig. 1b, lane 5) or Ty3 intasome buffer (Fig. 1b, lane 6). After each binding step, samples were collected and analysed by electrophoretic mobility shift assays (EMSAs) in 1% agarose gels. Detection of the Cy5 and FITC fluorophores present in the LTR and target DNA, respectively, was performed using a Typhoon FLA 9000 gel imaging scanner (GE Healthcare). For cryo-EM analysis, the sample was concentrated to ∼400 μl, centrifuged and loaded into a Superose 6 increase 10/300 GL column (GE Healthcare) equilibrated with 40 mM Tris pH 7, 80 mM NaCl, 7 mM MgCl_2_ and 1 mM DTT (Extended Data Fig. 1c). Fractions corresponding to the first elution peak were pooled.

Assembly of the minimal integration machinery was performed in the absence of the TFIIIC transcription factor following the same protocol but using an unlabelled 55-bp target DNA (Extended Data Fig. 1d). Fractions corresponding to the first elution peak were pooled, concentrated to ∼0.01 mg/ml with an Amicon Ultra 0.5 30K NMWL (Thermo Fisher Sci.) centrifugal concentrator and used for subsequent EM analysis.

### Ty3 integration assay

Evaluation of the integration activity of the reconstituted Ty3 intasome was performed through analysis of the DNA products after targeting at the full RNA Polymerase III transcriptional machinery. Samples were collected after recruitment of TFIIIC, TFIIIB and Ty3 intasome to the t(GUC)K target DNA, which was Cy3- or FITC-labelled in the upstream or downstream terminus, respectively (Fig.1a). Reaction mixtures were prepared in the presence of Cy5-labelled or unlabelled intasomes and, as a control, the assay was also performed in the absence of integration machinery (Fig. 1c, d). In order to analyse the nucleic acid fraction, the protein content was degraded using the broad-spectrum serine protease proteinase K. Thus, the sample was mixed with a proteinase K mixture (2 mg/ml proteinase K, 2% SDS (w/v) and 200 mM EDTA pH 8) in a 10:1.1 volume ratio and incubated for 1.5 h at 37 °C. Then, the cleaved sample was loaded into a 4% agarose gel and identification of the integration products was performed through detection of the FITC, Cy5 and Cy3 fluorophores in a Typhoon FLA 9000 gel imaging scanner (GE Healthcare) (Fig. 1c, d). 50 bp DNA markers (N3236S, New England Biolabs) were used to identify the size of the resulting DNA products (Fig. 1e).

### Negative stain EM sample preparation, data collection and processing

Negative stain electron microscopy of the full integration complex (TFIIIB-TFIIIC-DNA- Intasome; 920 kDa), the minimal integration complex (TFIIIB-Intasome; 400 kDa) and the TFIIIB-TFIIIC-DNA complex (670 kDa) were performed after gel-filtration chromatography following the same protocol. In brief, 3 μl of sample were applied to glow discharged (1 min at 15 mA; PELCO EasiGlow) Quantifoil R 1.2/1.3 copper grids coated with a thin carbon film, incubated for 1 min, washed with ultra-filtered water and stained with 2% uranyl acetate for 30 sec. Data collection was performed in-house in a ThermoFisher Tecnai F20 TEM microscope operating at 200kV, equipped with a F416 CMOS camera (TVIPS) and at a 50,000x magnification corresponding to 1.732 Å per pixel. All data processing was carried out in Relion 3.0.4.

### Cryo-EM sample preparation and data collection

Cryo-EM samples of *S. cerevisiae* Ty3 intasome engaged with TFIIIB and the *tD(GUC)K* target DNA were prepared on Quantifoil R 1.2/1.3 (400 mesh) copper grids coated with a thin carbon film prepared in house. Grids were glow discharged for 30 s at 15 mA (PELCO EasyGlow) before the application of 2 μl of sample at ∼0.1 mg/ml. After 30 s incubation at 18 °C and 100% humidity, the grids were blotted (drain time: 0.5 s, blot force: 3, blot time: 3 s) and plunged frozen into liquid ethane using a Vitrobot Mark IV system (ThermoFisher).

Separate data collections were performed for un-tilted and 30°-tilted datasets at the Electron Bio-Imaging Centre (eBIC) on a Titan Krios transmission electron microscope (ThermoFisher) at 300 KeV. For the un-tilted data collection, 4974 movies were collected using a K2 Summit direct electron detector (Gatan, Inc.) at a 1.47 Å calibrated pixel size and a nominal magnification of 130,000x (Extended Data Fig. 2a). Fifty frames were collected per movie at a defocus range between -1.7 μm and -3.2 μm and a total exposure of 7 s (dose rate of 7.14 e^-^/ Å^2^/s, an accumulated total dose of ∼50 e^-^/ Å^2^ and a fractionated dose per frame of 1 e^-^/ Å^2^). For the 30°-tilted data collection, 2437 movies were collected in a K3 direct electron detector (Gatan, Inc.) at a nominal magnification of 81,000x and a calibrated pixel size of 0.53 Å (Extended Data Fig. 2b). Each movie was fractionated over 70 frames and collected for 5.3 s at a dose per frame of 1 e^-^/ Å^2^ and at a defocus range from -2.4 μm to -4.0 μm, which yielded a dose rate of 13.20 e^-^/ Å^2^/s and an accumulated total dose of ∼70 e^-^/ Å^2^.

### Cryo-EM Image Processing

Frame alignment and dose-weighting steps were performed on-the-fly during data collection using MotionCor2 software^70^ and CFFIND4^71^ was used for the estimation of the contrast transfer function (CTF) parameters. Unless stated otherwise, all the subsequent steps of particle picking, extraction, 2D and 3D classification, and post-processing were carried out in Relion (versions 3.0.4 to 3.1.1)^72^.

For the un-tilted dataset, 1,090,240 particles were auto-picked and subjected to three steps of classification that provided highly-detailed 2D class averages, which accounted for a subset of 386,471 particles. Then, the particles were imported into CryoSPARC 2.0^73^ and subjected to an *ab-initio* 3D classification process, which provided five major classes. Class #0 (∼86,939 particles) and class #4 (∼94,355 particles) showed the characteristic tetrameric organisation observed in other intasome reconstructions (i.e. PFV or HIV) and the presence of an extra unidentified density. After joining the particles from these two classes, a hetero-refinement step was carried out, which resulted in two new 3D reconstructions: class #0 (∼55,546 particles) at 9.65 Å-resolution and class #1 (∼125,748 particles) at 5.40 Å-resolution. Particles corresponding to class #1 were re-extracted and imported into Relion, where they were subjected to a refinement and post-processing step that resulted in a final 3.8 Å-resolution map, according to the gold-standard FSC cut-off criterion at 0.143. The resulting map showed unequivocally the presence of the Ty3 intasome complex (containing four molecules of Ty3 integrase and two LTR fragments) engaged with the target DNA fragment and the Brf1 and TBP transcription factors. However, detailed analysis of the map and of the particle orientation distribution sphere indicated the existence of preferential orientation, which caused the presence of anisotropic features in the reconstruction.

For the 30°-tilted dataset, 994,999 particles were auto-picked, extracted and subjected to five steps of 2D classification, which resulted in a subset of 242,527 particles. 3D classification of this subset provided three classes that showed anisotropic characteristics similar to those observed previously in the un-tilted dataset.

With the aim of solving these problems, the un-tilted and 30°-tilted datasets were merged following a protocol described elsewhere^74^. In brief, after calculating the “real” pixel size from the obtained 3D reconstructions, particles from the un-tilted 2D- classification subset were re-extracted and re-sized to the nominal pixel size of 1.06 Å observed in the tilted dataset. Then, the scaled particles were merged with the 2D- classification subset of the tilted data collection, resulting in a new dataset of 628,998 particles (Extended Data Fig. 2c). Next, 3D classification was performed using a 20 Å-filtered map obtained from the un-tilted dataset, which resulted in five different classes (Extended Data Fig. 2d). Class #3 (169,140 particles), Class #4 (116,343 particles) and Class #5 (118,081 particles) showed density features corresponding to Ty3 intasome and TFIIIB. After joining of the particles, a 3D refinement was performed, which rendered a map at 3.97 Å-resolution, according to the gold-standard FSC cut- off criterion at 0.143. Then, to improve the quality around specific regions of the map, we performed a hierarchical 3D classification process without alignment using masks around the intasome, TFIIIB and the peripheral CTD-CHD domains (Extended Data Fig. 2d). The resulting class was subjected to a CTF refinement process that encompassed trefoil and 4^th^ order aberration corrections, followed by correction of magnification anisotropy and finally, defocus refinement on a per particle basis to correct CTF estimation errors for the titled particles. Next, a symmetry expansion process followed by further consensus refinement and post-processing steps rendered a map which showed improved features and density corresponding to side- chains. Local resolution analysis reported a global resolution of 3.98 Å-resolution at the gold-standard 0.143 FSC cut-off criterion (Extended Data Fig. 2e-g).

### Cryo-EM Model Building and Refinement

An initial model of Ty3 integrase was obtained using the Phyre2 modelling web portal^75^, which reported structural similarity to the prototype foamy virus (PFV) integrase. Four copies of the predicted Ty3 integrase were structurally aligned to the tetrameric structure of PFV^76^ (RCSB Protein Data Bank (PDB) code: 3OS0), which rendered a preliminary *in-silico* model of the Ty3 intasome. A rigid-body fit of the model into the cryo-EM map was subsequently performed in UCSF Chimera (version 1.14)^77^, which provided the estimated position and general orientation of Ty3 intasome. In order to account for differences in the relative orientation of the Ty3 integrase domains, each subunit was split into its constituting domains: N-terminal extended domain (NED), N-terminal domain (NTD), catalytic core domain (CCD) and C-terminal domains (CTD). The resulting individual domains were rigidly-fitted into the cryo-EM map in Chimera, which led to an improved positioning of the subunits. Manual correction of the connecting regions was carried out in Coot (version 0.8.9.2)^78^ and it was guided by secondary structure prediction performed in the PsiPred webserver (http://bioinf.cs.ucl.ac.uk/psipred/). Following manual fitting, the model was subjected to real-space refinement in the Phenix Suite (version 1.18.1-3865)^79^ and the process was repeated iteratively until a tetrameric model of Ty3 intasome was obtained.

At this stage, regions of unassigned density were still observed in the map and we proceeded to the rigid-body fit of the TFIIIB transcription factor (PDB code: 6EU0) into them using Chimera. We carried out a manual and automatic refinement in Coot and Phenix, as previously described, which allowed us to build the majority of the Brf1 and TBP models. Density corresponding to Bdp1 protein was very poor and only a rigid-body fit of the SANT domain was performed using Chimera.

Next, DNA fragments of the PFV gene LTRs (PDB code: 3OS0) were used as a reference model and rigidly fitted into the map using Chimera. Following *in-silico* mutagenesis to correct the sequence discrepancies, the Ty3 LTRs were real-space refined in Phenix. Similarly, model building of the target DNA was based on structural alignment of TFIIIB bound to the *SNR6* promoter (PDB code: 6EU0), which allowed us to determine the position of the TATA-box element. Nucleotides were then mutated into the *tD(GUC)K* promoter sequence. Next, the target DNA model was extended by an iterative process consisting of the manual addition of 3-4 nucleotides at the downstream end and its refinement in Phenix. This method was repeated recursively until the DNA molecule near the intasome active site was built. Here, the structure of HIV strand transfer complex^46^ (PDB code: 5U1C) was used as a reference to determine the position of the LTR reactive strands and to build the 5-nt target site duplication (TSD), which is characteristic of Ty3 intasome. Previous studies into the local features of Ty3 targeting^48^ were taken into account in order to ensure the nucleotide registry around the integration sites was correctly modelled.

After model building of TFIIIB, Ty3 intasome and the target DNA, we observed the presence of two extra densities in the cryo-EM map: one close to the TFIIIB transcription factor and another near the DNA downstream end. After building of the Ty3 intasome, the C-terminal domain (CTD) of a peripheral integrase subunit remained unmodelled, since its position differed from that observed in similar structures. However, detailed analysis of the unassigned densities showed that the missing Ty3 CTD could be rigidly-fit with high confidence into the extra map near TFIIIB. Following docking, the linker to the catalytic-core domain (CCD) was built using Coot and this region was subjected to real-space refinement in Phenix. After this process, part of the extra map near Brf1 still remained unidentified. This density was connected to the newly built CTD and it seemed to correspond to the very C-terminus of the Ty3 integrase, a region absent in retroviral homologs. In order to determine the protein architecture of this region (aa. 440-504), an intensive *in-silico* modelling was carried out in Phyre2 which predicted its folding into a chromo-domain (CHD). The generated Ty3 CHD was accurately fitted into the map near Brf1 using Chimera, and refined in Phenix after manual correction in Coot.

Finally, the remaining unassigned density near the downstream DNA region was also situated close to the C-terminal domain of a core integrase subunit, which prompted us to consider whether this feature also corresponded to a chromo-domain. Albeit the map quality around this area was poor, rigid-body fit of the CHD model into the unsharpened cryo-EM map unequivocally identified this density as a chromo-domain fold.

Cryo-EM figures were prepared using UCSF Chimera 1.14 and ChimeraX-1.0. Multiple sequence alignments were calculated using the Clustalw Omega server (https://www.ebi.ac.uk/Tools/msa/clustalo/) and figures were prepared in Jalview 2.10.1. Structural comparisons were performed in Pymol from the corresponding PDB codes obtained in the RCSB database (https://www.rcsb.org/).

## DATA AVAILABILITY

The structure of the Ty3 intasome-TFIIIB-tRNA promoter complex and its associated data have been deposited into the Protein Data Bank under accession code XXXX, and the Electron Microscopy Data Bank under accession codes XXXX.

## ACKNOWLEDGEMENTS

We thank all the members of the Vannini lab for critically reading the manuscript and for fruitful discussions. We further acknowledge the eBIC Cryo-EM facilities at the UK national electron bio-imaging centre at Diamond, funded by the Wellcome Trust, MRC and BBSRC, for access and support under proposal BI21809. This work was funded by the Cancer Research UK Programme Foundation (CR-UK C47547/A21536) and a Wellcome Trust Investigator Award (200818/Z/16/Z) to A.V.

## AUTHOR CONTRIBUTIONS

G.A.-P. designed and performed the experiments, analysed the data and wrote the manuscript. L.J. produced recombinant TFIIIC. C.P.-P. cloned and carried out initial characterization of Ty3 integrase. F.B. carried out cryo-EM sample preparation, screening and sample collection and helped with data analysis. A.V. designed and coordinated the project, analysed data and wrote the manuscript.

---

**Extended Data Figure 1.**
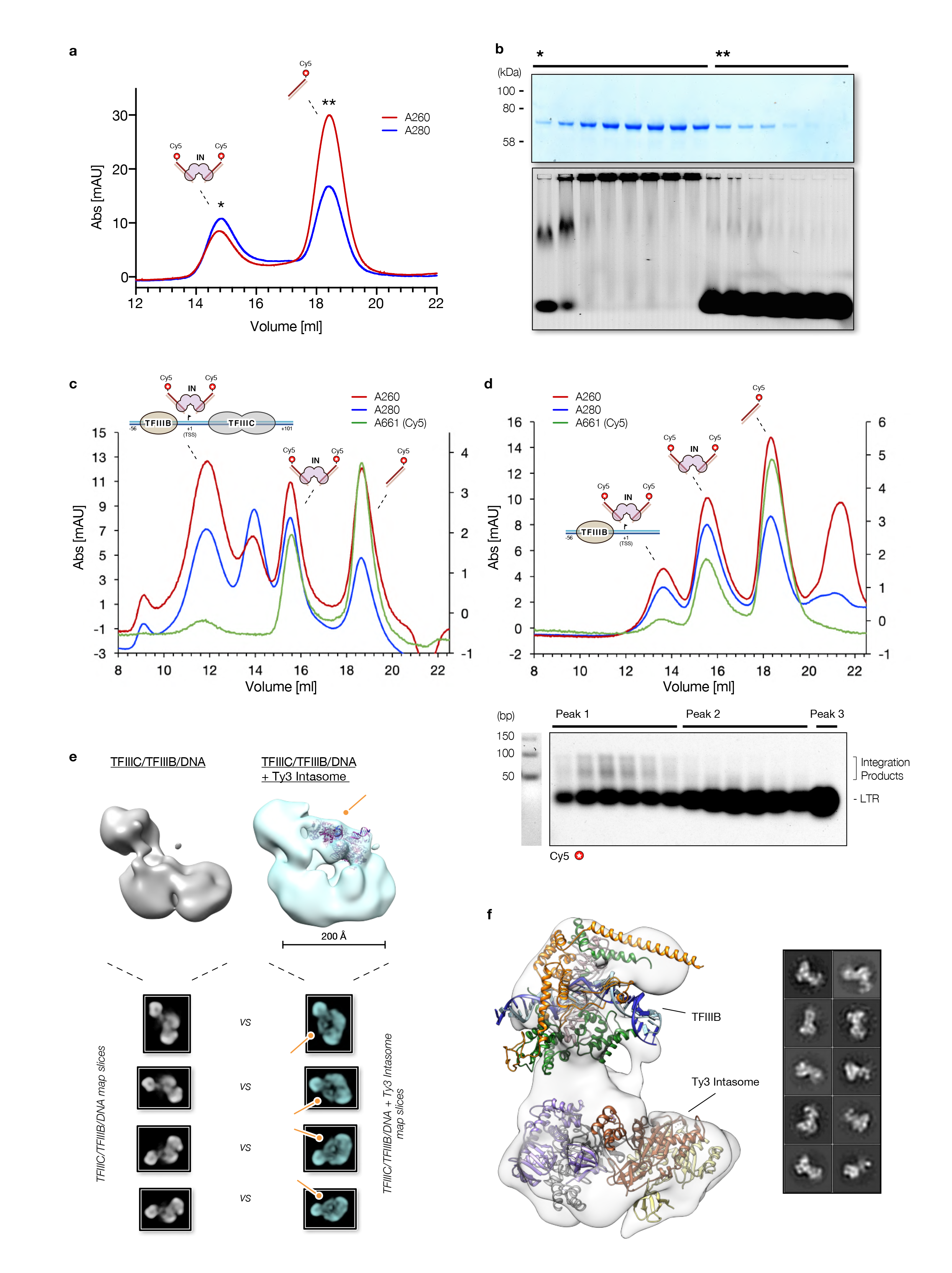
Reconstitution of Ty3 integration machinery and targeting complexes. **a,** Gel-filtration chromatography of Ty3 intasome. UV absorbance at 260 and 280 nm are indicated as red and blue lines, respectively. **b,** Analysis of the gel-filtration fractions by SDS-PAGE *(top)* showed the presence of Ty3 integrase only in the first elution peak. Analysis in 4.5% agarose gels *(bottom)* found free LTR in the second peak and the formation of a higher order complex in the first elution peak, suggesting the presence of Ty3 intasome in these fractions. **c,** Gel-filtration chromatography of full Ty3 integration machinery (TFIIIC/TFIIIB/DNA/Intasome). **d,** Gel-filtration chromatography of minimal Ty3 integration machinery (TFIIIB/DNA/Intasome). DNA integration products were analysed in 4.5% agarose gels after proteinase K treatment (*inset)*. **e,** Negative stain electron microscopy analysis of the TFIIIC/TFIIIB/DNA/Intasome complex. Comparison of EM reconstructions obtained in the absence *(left)* and presence *(right)* of Ty3 intasome shows the existence of an extra density on the periphery of the complex. Fitting of a homology model of Ty3 intasome (ribbon, 240 kDa) into the extra density shows a size compatible with a Ty3 integrase tetramer. Central slices of the negative stain maps *(insets)* also confirm the presence of the extra region (orange arrows). **f,** Negative stain reconstruction of the minimal integration machinery. *S. cerevisiae* TFIIIB and a homology model of Ty3 integrase (four subunits) are fitted in the map. Representative 2D class-averages are also shown *(inset)*.

**Extended Data Figure 2.**
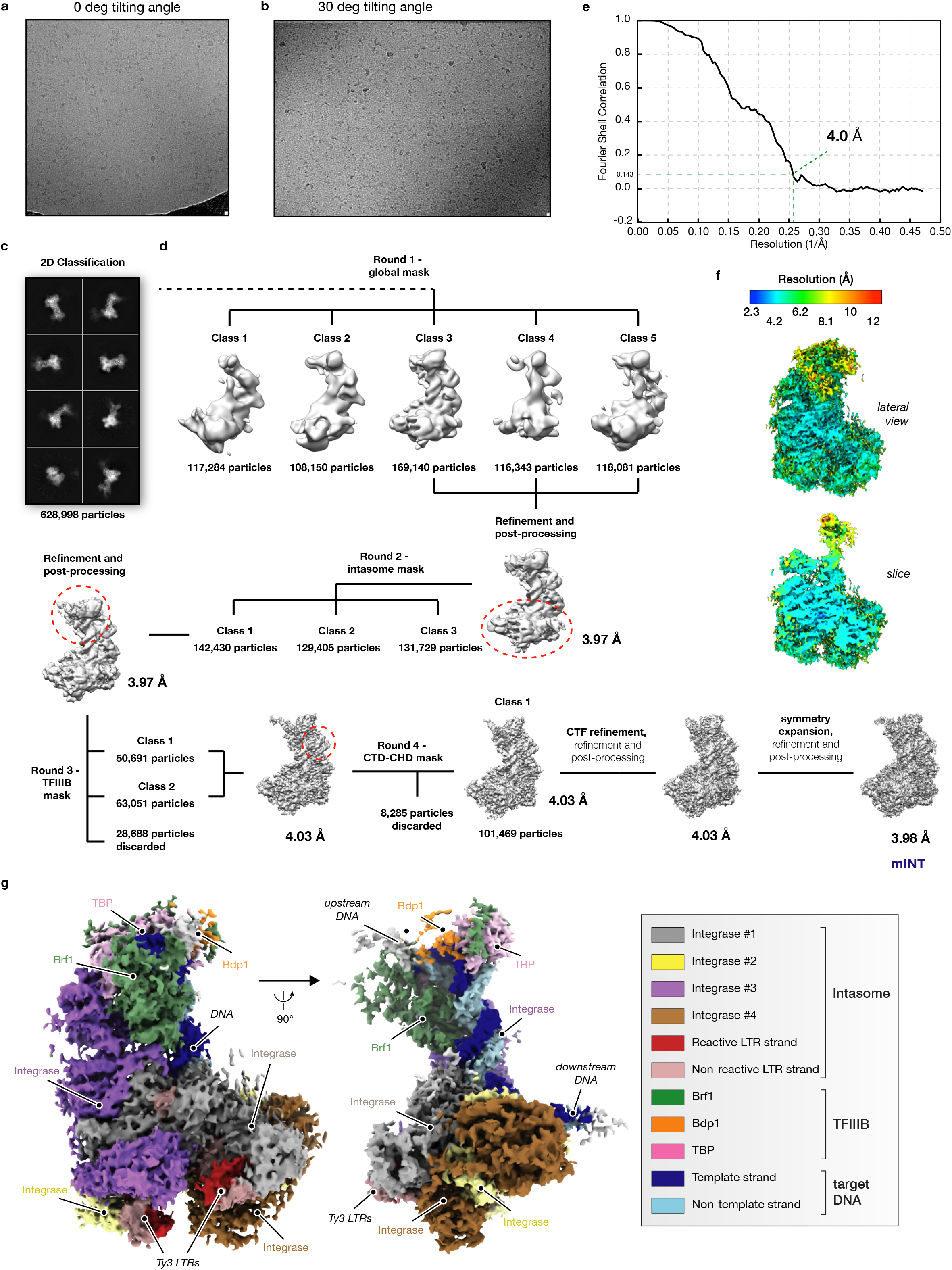
Cryo-EM data processing, reconstruction and resolution estimation a, b, Representative raw micrographs of TFIIIB - Ty3 Intasome datasets collected at 0° tilting angle (a) and 30° tilting angle (b). **c,** Eight reference-free 2D class averages. **d,** 3D classification of the joined set of particles from 0° and 30° tilting angles datasets. The particles were subjected to a hierarchical process, encompassing several rounds of classification using global or focused masks (dashed red circles around specific regions of the complex), as described in the schematic. The estimated resolution at the gold-standard FSC (FSC= 0.143) and the number of particles contributing to each class are indicated close to the corresponding 3D reconstructions. **e,** Fourier-shell correlation (FSC) representation of the cryo-EM reconstruction with the estimated resolution at the gold-standard FSC. **f,** Resolution estimation of the cryo-EM map calculated with ResMap. Lateral *(top)* and central slice *(bottom)* views are shown and coloured according to the local resolution, as indicated in the scale bar. **g,** Cryo-EM reconstruction of the Ty3 retrotransposon targeting TFIIIB bound to *tD(GUC)K* DNA promoter. TFIIIB subunits, DNA molecules and Ty3 intasome monomers are coloured as indicated in the table.

**Extended Data Figure 3.**
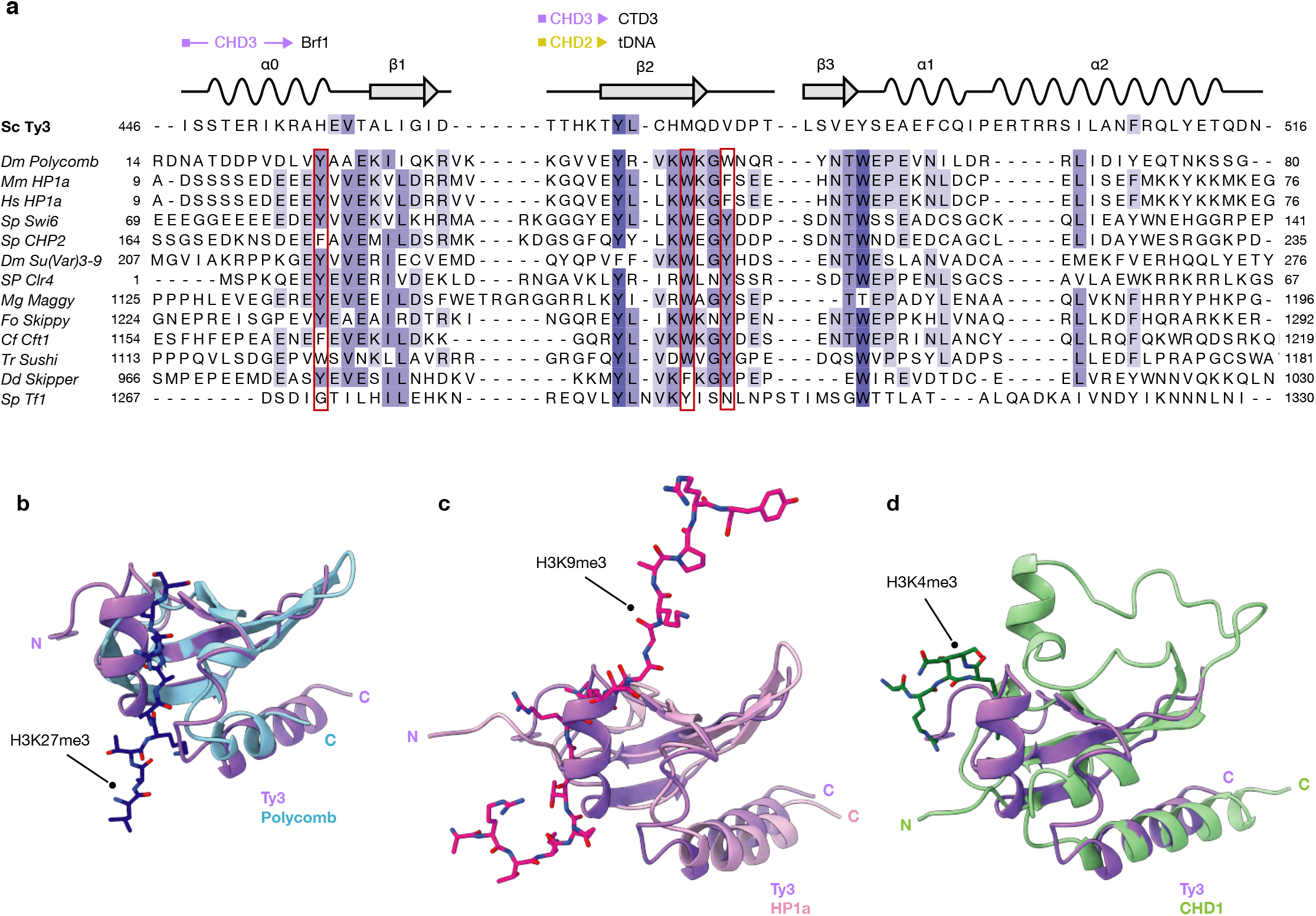
Chromodomain sequence and structure comparison. **a,** Sequence alignment of representative chromodomains from different species (Abbreviations: Sc, *Saccharomyces. cerevisiae*; Dm, *Drosophila melanogaster*; Hs, *Homo sapiens*; Sp, *Schizosaccharomyces pombe*; Mg, *Magnaporthe grisea*; Fo, *Fusarium oxysporum*; Cf, *Cladosporium fulvum*; Tr, *Takifugu rubripes;* Dd, *Dictyostelium discoideum*). The aromatic cage residues required for recognition of histone tail peptides are outlined with a red box. The schematic representation of Ty3 CHD motif organisation is shown above the sequence alignment. The regions of Ty3 CHD involved in interactions with other integrase domains or factors are delimited by an arrowed line, coloured and labelled according to the chromodomain subunit mediating the binding, as in Fig. 2c. Structure comparison of *S. cerevisiae* Ty3 integrase chromodomain (purple, this work) and **b,** *D. melanogaster* Polycomb chromodomain (light blue) complexed with H3K27me3 histone tail (dark blue) (PDB code: 1PDQ); **c,** *M. musculus* HP1a chromodomain (light pink) bound to the H3K9me3 histone tail (dark pink) (PDB code: 2RVN) and **d,** *H. sapiens* CHD1 first chromodomain (light green) complexed with H3K4me3 (dark green) (PDB code: 2B2W).

**Extended Data Figure 4.**
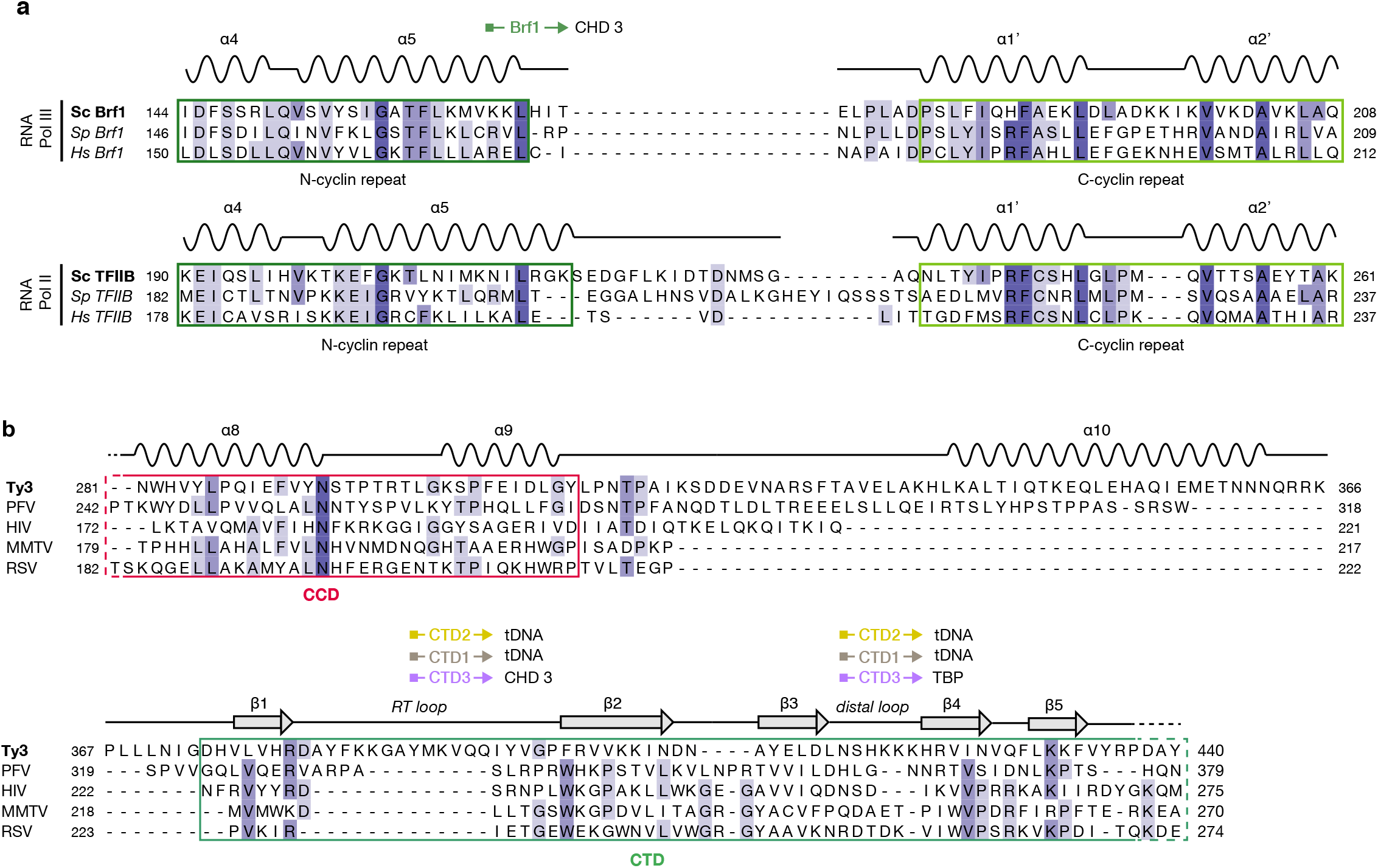
Sequence comparisons. **a,** Multiple sequence alignments of the linker region between the B-core cyclin-repeats of Brf1 *(up)* and TFIIB *(down)* transcription factors in *S. cerevisiae*, *S. pombe* and *H. sapiens*. Secondary structure motifs are represented above each alignment. **b,** Multiple sequence alignment of the integrase CCD-CTD region of *S. cerevisiae* Ty3 and PFV, HIV, MMTV and RSV retroviruses. Secondary structure representation shows the position of the conserved motifs. Catalytic core domains (CCD) and C-terminal domains (CTD) are delimited by red and green boxes, respectively. The regions of Ty3 retrotransposon involved in interactions with other integrase domains or factors are delimited by an arrowed line, coloured and labelled according to the specific integrase subunit mediating the binding, as in Fig 2c.

## REFERENCES

1. I. Ahmadi, A., De Toma, I., Vilor-Tejedor, N., Eftekhariyan Ghamsari, M. R. & Sadeghi, Transposable elements in brain health and disease. Ageing Res Rev 64, 101153, doi:10.1016/j.arr.2020.101153 (2020).

2. Burns, K. H. Transposable elements in cancer. Nat Rev Cancer 17, 415–424, doi:10.1038/nrc.2017.35 (2017).

3. Nowak, E. et al. Ty3 reverse transcriptase complexed with an RNA-DNA hybrid shows structural and functional asymmetry. Nat Struct Mol Biol 21, 389–396, doi:10.1038/nsmb.2785 (2014).

4. Andrake, M. D. & Skalka, A. M. Retroviral Integrase: Then and Now. Annu Rev Virol 2, 241–264, doi:10.1146/annurev-virology-100114-055043 (2015).

5. Lesbats, P., Engelman, A. N. & Cherepanov, P. Retroviral DNA Integration. Chem Rev 116, 12730–12757, doi:10.1021/acs.chemrev.6b00125 (2016).

6. Chalker, D. L.,, Sandmeyer, S. B. & Sites of RNA polymerase III transcription initiation and Ty3 integration at the U6 gene are positioned by the TATA box. 90, 4927–4931 (1993).

7. Sandmeyer, S., Patterson, K. & Bilanchone, V. Ty3, a Position-specific Retrotransposon in Budding Yeast. Microbiol Spectr 3, MDNA3-0057-2014, doi:10.1128/microbiolspec.MDNA3-0057-2014 (2015).

8. Leem, Y. E. et al. Retrotransposon Tf1 is targeted to Pol II promoters by transcription activators. Mol Cell 30, 98–107, doi:10.1016/j.molcel.2008.02.016 (2008).

9. Levin, H. L. & Moran, J. V. Dynamic interactions between transposable elements and their hosts. Nat Rev Genet 12, 615–627, doi:10.1038/nrg3030 (2011).

10. Sultana, T., Zamborlini, A., Cristofari, G. & Lesage, P. Integration site selection by retroviruses and transposable elements in eukaryotes. Nat Rev Genet 18, 292–308, doi:10.1038/nrg.2017.7 (2017).

11. Xie, W. et al. Targeting of the yeast Ty5 retrotransposon to silent chromatin is mediated by interactions between integrase and Sir4p. Mol Cell Biol 21, 6606–6614, doi:10.1128/MCB.21.19.6606-6614.2001 (2001).

12. Bachman, N., Gelbart, M. E., Tsukiyama, T. & Boeke, J. D. TFIIIB subunit Bdp1p is required for periodic integration of the Ty1 retrotransposon and targeting of Isw2p to S. cerevisiae tDNAs. Genes Dev 19, 955–964, doi:10.1101/gad.1299105 (2005).

13. Bridier-Nahmias, A., et al. Retrotransposons. An RNA polymerase III subunit determines sites of retrotransposon integration. Science 348, 585–588, doi:10.1126/science.1259114 (2015).

14. Cheung, S. et al. Ty1 Integrase Interacts with RNA Polymerase III-specific Subcomplexes to Promote Insertion of Ty1 Elements Upstream of Polymerase (Pol) III-transcribed Genes. J Biol Chem 291, 6396–6411, doi:10.1074/jbc.M115.686840 (2016).

15. Curcio, M. J., Lutz, S. & Lesage, P. The Ty1 LTR-Retrotransposon of Budding Yeast, Saccharomyces cerevisiae. Microbiol Spectr 3, MDNA3-0053-2014, doi:10.1128/microbiolspec.MDNA3-0053-2014 (2015).

16. Cherepanov, P., Ambrosio, A. L., Rahman, S., Ellenberger, T. & Engelman, A. Structural basis for the recognition between HIV-1 integrase and transcriptional coactivator p75. Proc Natl Acad Sci U S A 102, 17308–17313, doi:10.1073/pnas.0506924102 (2005).

17. Emiliani, S. et al. Integrase mutants defective for interaction with LEDGF/p75 are impaired in chromosome tethering and HIV-1 replication. J Biol Chem 280, 25517–25523, doi:10.1074/jbc.M501378200 (2005).

18. Turlure, F., Devroe, E., Silver, P. A. & Engelman, A. Human cell proteins and human immunodeficiency virus DNA integration. Front Biosci 9, 3187–3208, doi:10.2741/1472 (2004).

19. Cramer, P. et al. Structure of eukaryotic RNA polymerases. Annu Rev Biophys 37, 337–352, doi:10.1146/annurev.biophys.37.032807.130008 (2008).

20. Teichmann, M., Wang, Z. & Roeder, R. G. A stable complex of a novel transcription factor IIB- related factor, human TFIIIB50, and associated proteins mediate selective transcription by RNA polymerase III of genes with upstream promoter elements. Proc Natl Acad Sci U S A 97, 14200–14205, doi:10.1073/pnas.97.26.14200 (2000).

21. Abascal-Palacios, G., Ramsay, E. P., Beuron, F., Morris, E. & Vannini, A. Structural basis of RNA polymerase III transcription initiation. Nature 553, 301–306, doi:10.1038/nature25441 (2018).

22. Han, Y., Yan, C., Fishbain, S., Ivanov, I. & He, Y. Structural visualization of RNA polymerase III transcription machineries. Cell Discov 4, 40, doi:10.1038/s41421-018-0044-z (2018).

23. Kassavetis, G. A., Braun, B. R., Nguyen, L. H. & Geiduschek, E. P. S. cerevisiae TFIIIB is the transcription initiation factor proper of RNA polymerase III, while TFIIIA and TFIIIC are assembly factors. Cell 60, 235–245, doi:10.1016/0092-8674(90)90739-2 (1990).

24. Kassavetis, G. A., Letts, G. A. & Geiduschek, E. P. A minimal RNA polymerase III transcription system. EMBO J 18, 5042–5051, doi:10.1093/emboj/18.18.5042 (1999).

25. Vorlander, M. K., Khatter, H., Wetzel, R., Hagen, W. J. H. & Muller, C. W. Molecular mechanism of promoter opening by RNA polymerase III. Nature 553, 295–300, doi:10.1038/nature25440 (2018).

26. Moqtaderi, Z. et al. Genomic binding profiles of functionally distinct RNA polymerase III transcription complexes in human cells. Nat Struct Mol Biol 17, 635–640, doi:10.1038/nsmb.1794 (2010).

27. Vannini, A. & Cramer, P. Conservation between the RNA polymerase I, II, and III transcription initiation machineries. Mol Cell 45, 439–446, doi:10.1016/j.molcel.2012.01.023 (2012).

28. Ishiguro, A., Kassavetis, G. A. & Geiduschek, E. P. Essential roles of Bdp1, a subunit of RNA polymerase III initiation factor TFIIIB, in transcription and tRNA processing. Mol Cell Biol 22, 3264–3275, doi:10.1128/MCB.22.10.3264-3275.2002 (2002).

29. Chaussivert, N., Conesa, C., Shaaban, S. & Sentenac, A. Complex interactions between yeast TFIIIB and TFIIIC. J Biol Chem 270, 15353–15358, doi:10.1074/jbc.270.25.15353 (1995).

30. Kirchner, J., Connolly, C. M. & Sandmeyer, S. B. Requirement of RNA polymerase III transcription factors for in vitro position-specific integration of a retroviruslike element. Science 267, 1488–1491, doi:10.1126/science.7878467 (1995).

31. Yieh, L., Hatzis, H., Kassavetis, G. & Sandmeyer, S. B. Mutational analysis of the transcription factor IIIB-DNA target of Ty3 retroelement integration. J Biol Chem 277, 25920–25928, doi:10.1074/jbc.M202729200 (2002).

32. Yieh, L., Kassavetis, G., Geiduschek, E. P. & Sandmeyer, S. B. The Brf and TATA-binding protein subunits of the RNA polymerase III transcription factor IIIB mediate position-specific integration of the gypsy-like element, Ty3. J Biol Chem 275, 29800–29807, doi:10.1074/jbc.M003149200 (2000).

33. Payer, L. M. & Burns, K. H. Transposable elements in human genetic disease. Nat Rev Genet 20, 760–772, doi:10.1038/s41576-019-0165-8 (2019).

34. Xue, B., Sechi, L. A. & Kelvin, D. J. Human Endogenous Retrovirus K (HML-2) in Health and Disease. Front Microbiol 11, 1690, doi:10.3389/fmicb.2020.01690 (2020).

35. Kury, P. et al. Human Endogenous Retroviruses in Neurological Diseases. Trends Mol Med 24, 379–394, doi:10.1016/j.molmed.2018.02.007 (2018).

36. Li, W. et al. Human endogenous retrovirus-K contributes to motor neuron disease. Sci Transl Med 7, 307ra153, doi:10.1126/scitranslmed.aac8201 (2015).

37. Chen, J., Foroozesh, M. & Qin, Z. Transactivation of human endogenous retroviruses by tumor viruses and their functions in virus-associated malignancies. Oncogenesis 8, 6, doi:10.1038/s41389-018-0114-y (2019).

38. Salavatiha, Z., Soleimani-Jelodar, R. & Jalilvand, S. The role of endogenous retroviruses-K in human cancer. Rev Med Virol 30, 1–13, doi:10.1002/rmv.2142 (2020).

39. Lee, G. E., Mauro, E., Parissi, V., Shin, C. G. & Lesbats, P. Structural Insights on Retroviral DNA Integration: Learning from Foamy Viruses. Viruses 11, doi:10.3390/v11090770 (2019).

40. Li, X., Krishnan, L., Cherepanov, P. & Engelman, A. Structural biology of retroviral DNA integration. Virology 411, 194–205, doi:10.1016/j.virol.2010.12.008 (2011).

41. Maskell, D. P. et al. Structural basis for retroviral integration into nucleosomes. Nature 523, 366–369, doi:10.1038/nature14495 (2015).

42. Wilson, M. D. et al. Retroviral integration into nucleosomes through DNA looping and sliding along the histone octamer. Nat Commun 10, 4189, doi:10.1038/s41467-019-12007-w (2019).

43. Valkov, E. et al. Functional and structural characterization of the integrase from the prototype foamy virus. Nucleic Acids Res 37, 243–255, doi:10.1093/nar/gkn938 (2009).

44. Yin, Z. et al. Crystal structure of the Rous sarcoma virus intasome. Nature 530, 362–366, doi:10.1038/nature16950 (2016).

45. Ballandras-Colas, A. et al. Cryo-EM reveals a novel octameric integrase structure for betaretroviral intasome function. Nature 530, 358–361, doi:10.1038/nature16955 (2016).

46. Passos, D. O. et al. Cryo-EM structures and atomic model of the HIV-1 strand transfer complex intasome. Science 355, 89–92, doi:10.1126/science.aah5163 (2017).

47. Aye, M., Dildine, S. L., Claypool, J. A., Jourdain, S. & Sandmeyer, S. B. A truncation mutant of the 95-kilodalton subunit of transcription factor IIIC reveals asymmetry in Ty3 integration. Mol Cell Biol 21, 7839–7851, doi:10.1128/MCB.21.22.7839-7851.2001 (2001).

48. Patterson, K. et al. Local features determine Ty3 targeting frequency at RNA polymerase III transcription start sites. Genome Res 29, 1298–1309, doi:10.1101/gr.240861.118 (2019).

49. Butryn, A. et al. Structural basis for recognition and remodeling of the TBP:DNA:NC2 complex by Mot1. Elife 4, 1–22, doi:10.7554/eLife.07432 (2015).

50. Malik, H. S. & Eickbush, T. H. Modular evolution of the integrase domain in the Ty3/Gypsy class of LTR retrotransposons. J Virol 73, 5186–5190, doi:10.1128/JVI.73.6.5186-5190.1999 (1999).

51. Youkharibache, P. et al. The Small beta-Barrel Domain: A Survey-Based Structural Analysis. Structure 27, 6–26, doi:10.1016/j.str.2018.09.012 (2019).

52. Eissenberg, J. C. Structural biology of the chromodomain: form and function. Gene 496, 69–78, doi:10.1016/j.gene.2012.01.003 (2012).

53. Fischle, W. et al. Molecular basis for the discrimination of repressive methyl-lysine marks in histone H3 by Polycomb and HP1 chromodomains. Genes Dev 17, 1870–1881, doi:10.1101/gad.1110503 (2003).

54. Flanagan, J. F. et al. Double chromodomains cooperate to recognize the methylated histone H3 tail. Nature 438, 1181–1185, doi:10.1038/nature04290 (2005).

55. Shimojo, H. et al. Extended string-like binding of the phosphorylated HP1alpha N-terminal tail to the lysine 9-methylated histone H3 tail. Sci Rep 6, 22527, doi:10.1038/srep22527 (2016).

56. Hansen, L. J. & Sandmeyer, S. B. Characterization of a transpositionally active Ty3 element and identification of the Ty3 integrase protein. J Virol 64, 2599–2607, doi:10.1128/JVI.64.6.2599-2607.1990 (1990).

57. Chatterjee, A. G., Leem, Y. E., Kelly, F. D. & Levin, H. L. The chromodomain of Tf1 integrase promotes binding to cDNA and mediates target site selection. J Virol 83, 2675–2685, doi:10.1128/JVI.01588-08 (2009).

58. Gao, X., Hou, Y., Ebina, H., Levin, H. L. & Voytas, D. F. Chromodomains direct integration of retrotransposons to heterochromatin. Genome Res 18, 359–369, doi:10.1101/gr.7146408 (2008).

59. Christ, F. et al. Small-molecule inhibitors of the LEDGF/p75 binding site of integrase block HIV replication and modulate integrase multimerization. Antimicrob Agents Chemother 56, 4365–4374, doi:10.1128/AAC.00717-12 (2012).

60. Poeschla, E. M. Integrase, LEDGF/p75 and HIV replication. Cell Mol Life Sci 65, 1403–1424, doi:10.1007/s00018-008-7540-5 (2008).

61. Engelman, A. & Cherepanov, P. The lentiviral integrase binding protein LEDGF/p75 and HIV-1 replication. PLoS Pathog 4, e1000046, doi:10.1371/journal.ppat.1000046 (2008).

62. Dieci, G., Percudani, R., Giuliodori, S., Bottarelli, L. & Ottonello, S. TFIIIC- independent in vitro transcription of yeast tRNA genes. J Mol Biol 299, 601–613, doi:10.1006/jmbi.2000.3783 (2000).

63. Connolly, C. M. & Sandmeyer, S. B. RNA polymerase III interferes with Ty3 integration. FEBS Lett 405, 305–311, doi:10.1016/s0014-5793(97)00200-7 (1997).

64. Asif-Laidin, A. et al. A small targeting domain in Ty1 integrase is sufficient to direct retrotransposon integration upstream of tRNA genes. EMBO J 39, e104337, doi:10.15252/embj.2019104337 (2020).

65. Gerlach, V. L., Whitehall, S. K., Geiduschek, E. P. & Brow, D. A. TFIIIB placement on a yeast U6 RNA gene in vivo is directed primarily by TFIIIC rather than by sequence-specific DNA contacts. Mol Cell Biol 15, 1455–1466, doi:10.1128/MCB.15.3.1455 (1995).

66. Male, G. et al. Architecture of TFIIIC and its role in RNA polymerase III pre-initiation complex assembly. Nat Commun 6, 7387, doi:10.1038/ncomms8387 (2015).

67. Stillman, D. J. & Geiduschek, E. P. Differential binding of a S. cerevisiae RNA polymerase III transcription factor to two promoter segments of a tRNA gene. EMBO J 3, 847–853 (1984).

68. Weissmann, F. et al. biGBac enables rapid gene assembly for the expression of large multisubunit protein complexes. Proc Natl Acad Sci U S A 113, E2564–2569, doi:10.1073/pnas.1604935113 (2016).

69. Juo, Z. S., Kassavetis, G. A., Wang, J., Geiduschek, E. P. & Sigler, P. B. Crystal structure of a transcription factor IIIB core interface ternary complex. Nature 422, 534–539, doi:10.1038/nature01534 (2003).

70. Zheng, S. Q. et al. MotionCor2: anisotropic correction of beam-induced motion for improved cryo-electron microscopy. Nat Methods 14, 331–332, doi:10.1038/nmeth.4193 (2017).

71. Rohou, A. & Grigorieff, N. CTFFIND4: Fast and accurate defocus estimation from electron micrographs. J Struct Biol 192, 216–221, doi:10.1016/j.jsb.2015.08.008 (2015).

72. Scheres, S. H. RELION: implementation of a Bayesian approach to cryo-EM structure determination. J Struct Biol 180, 519–530, doi:10.1016/j.jsb.2012.09.006 (2012).

73. Punjani, A., Rubinstein, J. L., Fleet, D. J. & Brubaker, M. A. cryoSPARC: algorithms for rapid unsupervised cryo-EM structure determination. Nat Methods 14, 290–296, doi:10.1038/nmeth.4169 (2017).

74. Wilkinson, M. E., Kumar, A. & Casanal, A. Methods for merging data sets in electron cryo-microscopy. Acta Crystallogr D Struct Biol 75, 782–791, doi:10.1107/S2059798319010519 (2019).

75. Kelley, L. A., Mezulis, S., Yates, C. M., Wass, M. N. & Sternberg, M. J. The Phyre2 web portal for protein modeling, prediction and analysis. Nat Protoc 10, 845–858, doi:10.1038/nprot.2015.053 (2015).

76. Maertens, G. N., Hare, S. & Cherepanov, P. The mechanism of retroviral integration from X-ray structures of its key intermediates. Nature 468, 326–329, doi:10.1038/nature09517 (2010).

77. Pettersen, E. F. et al. UCSF Chimera--a visualization system for exploratory research and analysis. J Comput Chem 25, 1605–1612, doi:10.1002/jcc.20084 (2004).

78. Emsley, P. & Cowtan, K. Coot: model-building tools for molecular graphics. Acta Crystallogr D Biol Crystallogr 60, 2126–2132, doi:10.1107/S0907444904019158 (2004).

79. Afonine, P. V. et al. Real-space refinement in PHENIX for cryo-EM and crystallography. Acta Crystallogr D Struct Biol 74, 531–544, doi:10.1107/S2059798318006551 (2018).

